# Growth Rules for the Repair of Asynchronous Irregular Neuronal Networks after Peripheral Lesions

**DOI:** 10.1101/810846

**Authors:** Ankur Sinha, Christoph Metzner, Neil Davey, Roderick Adams, Michael Schmuker, Volker Steuber

## Abstract

Several homeostatic mechanisms enable the brain to maintain desired levels of neuronal activity. One of these, homeostatic structural plasticity, has been reported to restore activity in networks disrupted by peripheral lesions by altering their neuronal connectivity. While multiple lesion experiments have studied the changes in neurite morphology that underlie modifications of synapses in these networks, the underlying mechanisms that drive these changes are yet to be explained. Evidence suggests that neuronal activity modulates neurite morphology and may stimulate neurites to selective sprout or retract to restore network activity levels. We developed a new spiking network model, simulations of which accurately reproduce network rewiring after peripheral lesions as reported in experiments, to study these activity dependent growth regimes of neurites. To ensure that our simulations closely resemble the behaviour of networks in the brain, we deafferent a biologically realistic network model that exhibits low frequency Asynchronous Irregular (AI) activity as observed in cerebral cortex.

Our simulation results indicate that the re-establishment of activity in neurons both within and outside the deprived region, the Lesion Projection Zone (LPZ), requires opposite activity dependent growth rules for excitatory and inhibitory post-synaptic elements. Analysis of these growth regimes indicates that they also contribute to the maintenance of activity levels in individual neurons. Furthermore, in our model, the directional formation of synapses that is observed in experiments requires that pre-synaptic excitatory and inhibitory elements also follow opposite growth rules. Lastly, we observe that our proposed model of homeostatic structural plasticity and the inhibitory synaptic plasticity mechanism that also balances our AI network are both necessary for successful rewiring of the network.

**Author summary:** An accumulating body of evidence suggests that our brain can compensate for peripheral lesions by adaptive rewiring of its neuronal circuitry. The underlying process, structural plasticity, can modify the connectivity of neuronal networks in the brain, thus affecting their function. To better understand the mechanisms of structural plasticity in the brain, we have developed a novel model of peripheral lesions and the resulting activity-dependent rewiring in a simplified cortical network model that exhibits biologically realistic asynchronous irregular activity. In order to accurately reproduce the directionality and time course of rewiring after injury that is observed in peripheral lesion experiments, we derive activity dependent growth rules for different synaptic elements: dendritic and axonal contacts. Our simulation results suggest that excitatory and inhibitory synaptic elements have to react to changes in neuronal activity in opposite ways. We show that these rules result in a homeostatic stabilisation of activity in individual neurons. In our simulations, both synaptic and structural plasticity mechanisms are necessary for network repair. Furthermore, our simulations indicate that while activity is restored in neurons deprived by the peripheral lesion, the temporal firing characteristics of the network can be changed by the rewiring process.

## Introduction

Multiple plasticity mechanisms act simultaneously and at differing time scales on neuronal networks in the brain. Whilst synaptic plasticity is limited to the changes in efficacy of pre-existing synapses, *structural* plasticity includes the formation and removal of whole neurites and synapses. Thus, structural plasticity can cause major changes in network function through alterations in connectivity. Along with confirmation of structural plasticity in the adult brain [1–4], recent work has also shown that axonal boutons and branches [5–10], and both inhibitory [11, 12] and excitatory dendritic structures [13, 14] are highly dynamic even in physiological networks.

Stability in spite of such continuous plasticity requires homeostatic forms of structural plasticity. A multitude of lesion experiments provide evidence for homeostatic structural plasticity [15–26]. A common feature observed in these studies is the substantial network reorganisation that follows deafferentation. Recent time-lapse imaging studies of neurites in the cortex during the rewiring process show that both axonal [6, 10, 27] and dendritic structures display increased turnover rates [10, 13, 28, 29] in and around the area deafferented by the peripheral lesion, the LPZ. Specifically, while excitatory neurons outside the LPZ sprout new axonal collaterals into the LPZ, inhibitory neurons inside the LPZ extend new axons outwards [6]. Along with an increased excitatory dendritic spine gain [28] and a marked loss of inhibitory shaft synapses [11, 30] in the LPZ, the rewiring of synapses in the network successfully restores activity to deprived LPZ neurons in many cases.

Access to such data and recent advances in simulation technology have enabled computational modelling of activity dependent structural plasticity [31–37]. In their seminal work, Butz and van Ooyen introduced the Model of Structural Plasticity (MSP) framework [31]. They demonstrated its utility by simulating a peripheral lesioning study to explore the activity dependent growth rules of neurites [33, 34]. Their analysis suggests that the restoration of activity could only be caused by the experimentally noted increase in excitatory lateral projections into the LPZ if dendritic elements sprouted at a lower level of activity than their axonal counterparts. The MSP framework has since been partially implemented in the NEST simulator [38] and is an important tool for the computational modelling of structural plasticity [39, 40].

While investigating the capacity of simplified cortical balanced AI spiking neural networks [41] to store and recall associative memories [42], we wondered how deafferentation and subsequent connectivity updates that accompany the network repair process would affect its performance. Since the peripheral lesion model proposed by Butz and van Ooyen [33] was not based on a balanced cortical network model with biologically realistic AI activity, their hypothesised growth rules did not elicit repair in our simulations. Additionally, while providing salient testable predictions, the original MSP growth rules have specifically been developed for excitatory neurites only—they do not provide activity dependent growth rules for inhibitory neurites, nor do they reproduce the experimentally observed outgrowth of inhibitory axons from the LPZ. A complete, general computational model of peripheral lesioning in cortical networks is therefore still lacking.

Here, we present a novel computational model of peripheral lesioning and recovery in a simplified cortical spiking neural network with biologically realistic characteristics. In its physiological state, our network model is balanced by inhibitory Spike Timing Dependent Plasticity (STDP) so that it exhibits a low frequency AI spiking state similar to the mammalian cortex [41]. By deafferenting this network and reproducing a course of repair as reported in experimental work, we derive new independent activity dependent growth rules for all neurites—excitatory and inhibitory, pre-synaptic and post-synaptic.These growth rules result in the ingrowth of excitatory projections into and the outgrowth of inhibitory projections from the deafferented area that is observed in experiments. Although deduced from network simulations, we find that our growth rules also contribute to the stability of individual neurons by re-establishing their balance between excitation and inhibition (E-I balance). Furthermore, we show that both homeostatic processes in our model—synaptic plasticity and structural plasticity—are necessary for repair. Our model provides a new platform to study the structural and functional consequences of peripheral lesions in cortical networks.

## Results

### A new model of recovery in simplified cortical AI networks after peripheral lesions

Our network model consists of excitatory (E) and inhibitory (I) conductance based point neuron populations [43] distributed in a continuous two-dimensional toroidal grid. Neurons in the network are connected via synapses to simulate a simplified cortical AI network balanced by inhibitory STDP [41] (Fig. 1A). Apart from inhibitory synapses projecting from the inhibitory neurons to the excitatory ones (IE synapses), whose weights are modified by Vogels-Sprekeler symmetric inhibitory STDP, all synaptic conductances (II, EI, EE) are static. Structural plasticity, however, acts on all synapses in the network. We simulate a peripheral lesion in the balanced network by deafferenting a spatial selection of neurons to form the LPZ. For easier analysis, and as often done in experimental lesion studies, we divide the neuronal population into four regions relative to the LPZ (Fig. 1B).

**Fig 1.**
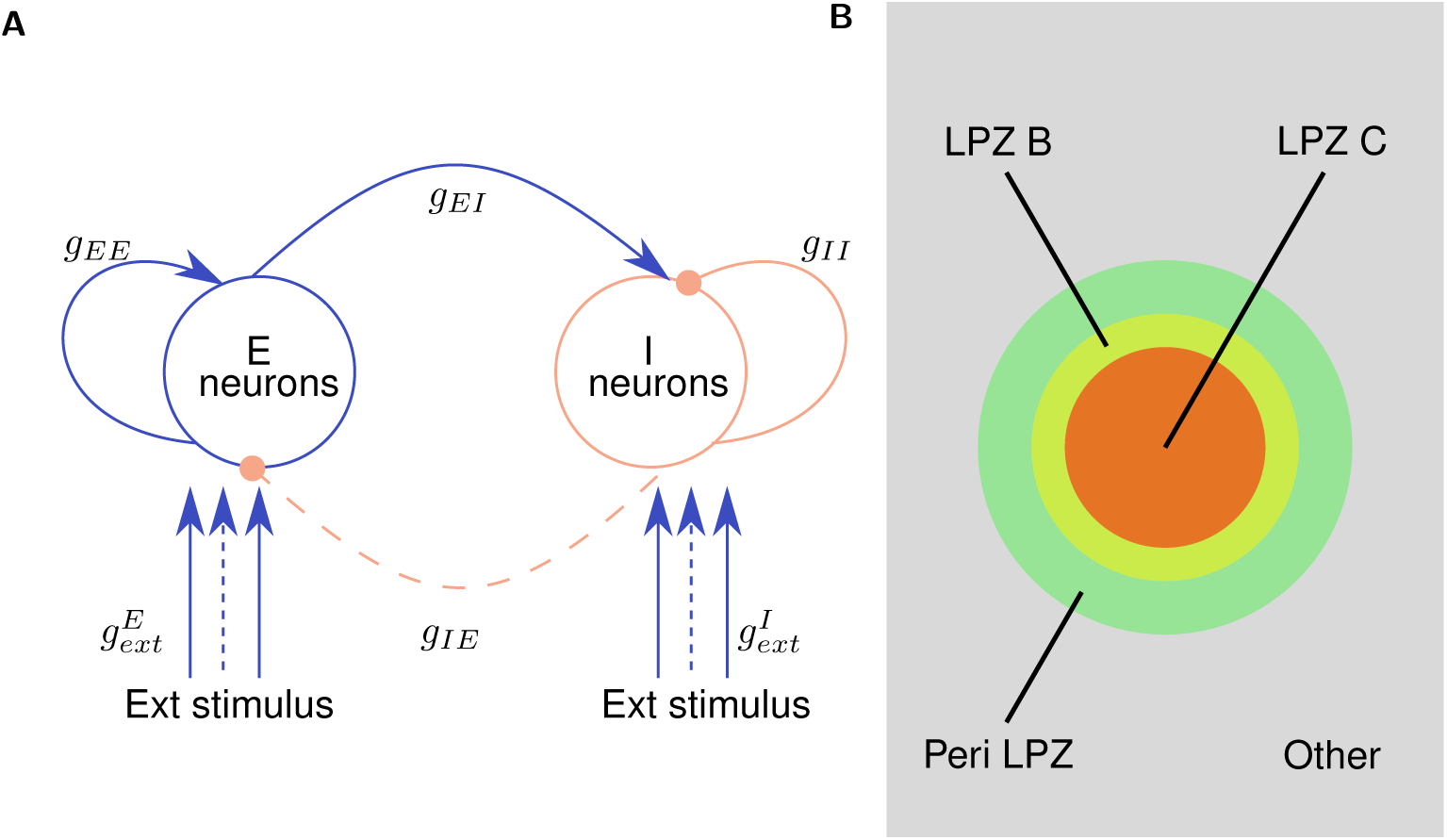
Overview of the model: **(A)** Excitatory (E) and Inhibitory (I) neurons (*N*_*E*_ = 4*N*_*I*_ (see Table 6)) are initially connected via synapses with a connection probability of (*p* = 0.02). All synapses (EE, EI, II), other than IE synapses, which are modulated by inhibitory spike-timing dependent plasticity, are static with conductances *g*_*EE*_, *g*_*EI*_, *g*_*II*_, respectively. All synapse sets are modifiable by the structural plasticity mechanism. External Poisson spike stimuli are provided to all excitatory and inhibitory neurons via static synapses with conductances 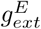 and 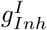, respectively. To simulate deafferentation, the subset of these synapses that project onto neurons in the Lesion Projection Zone (LPZ) (represented by dashed lines in the figure) are disconnected. **(B)** Spatial classification of neurons in relation to the LPZ: LPZ C (centre of LPZ) consists of 2.5 % of the neuronal population; LPZ B (inner border of LPZ) consists of 2.5 % of the neuronal population; Peri-LPZ (outer border of LPZ) consists of 5 % of the neuronal population; Other neurons consist of the remaining 90 % of the neuronal population. (Figure not to scale)

As in Butz and van Ooyen’s MSP framework, each neuron possesses sets of both pre-synaptic (axonal) and post-synaptic (dendritic) synaptic elements, the total numbers of which are represented by (*z*_*pre*_) and (*z*_*post*_), respectively. Excitatory and inhibitory neurons only possess excitatory 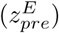 and inhibitory axonal elements 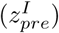, respectively, but they can each host both excitatory and inhibitory dendritic elements 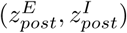 (Fig. 2A). The rate of change of each type of synaptic element, (*dz/dt*), is modelled as a Gaussian function of the neuron’s “calcium concentration” ([*Ca*^2+^]):

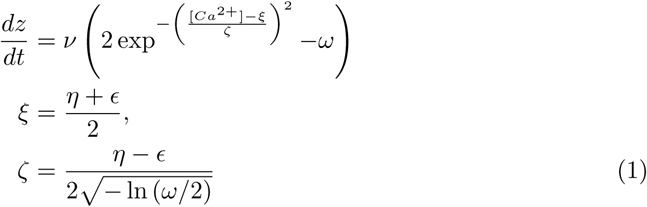

**Fig 2.**
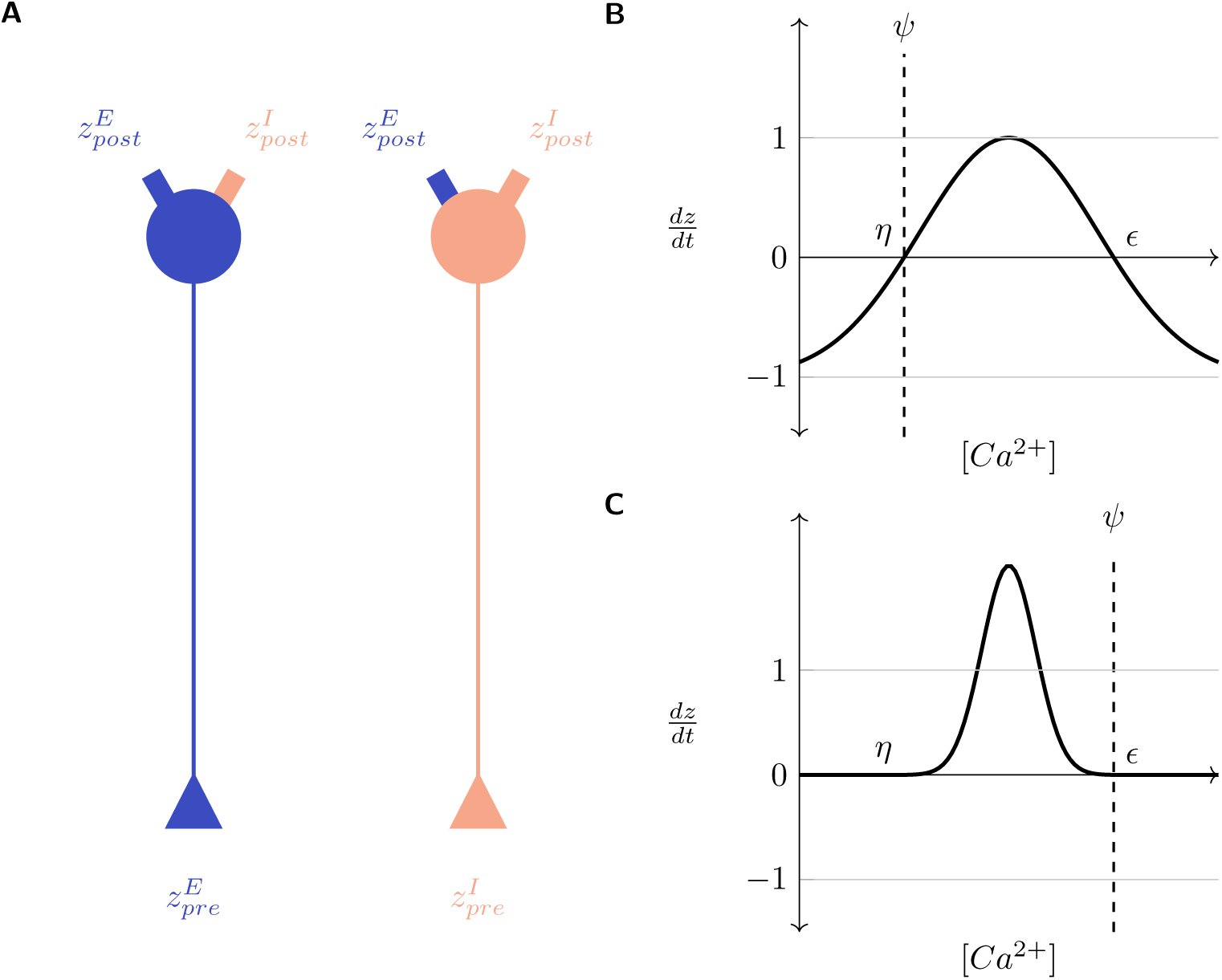
Gaussian growth curves modulate the rate of turnover of synaptic elements 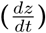 in a neuron as a function of its [*Ca*^2+^]: **(A)** Excitatory: Blue; Inhibitory: Red; All neurons possess excitatory and inhibitory post-synaptic elements 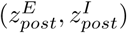 but excitatory and inhibitory neurons can only bear excitatory and inhibitory pre-synaptic elements, respectively 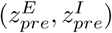; **(B)** and **(C)**: Example Gaussian growth curves. Constants *η* and control the width and positioning of the growth curve on the x-axis. *ω* (see Equation 1) controls the positioning of the growth curve on the y-axis. *ν* (see Equation 1) is a scaling factor. *ψ* is the optimal [*Ca*^2+^] for the neuron. The minimum and maximum values of *dz/dt* can be analytically deduced to be −*νω* and *ν*(2 − *ω*) respectively (see Methods). The relationship between *η*, *ϵ*, and *ψ* regulates the activity dependent dynamics of neurites. **(B)** *ψ* = *η* = 5.0, *ϵ* = 15.0, *ν* = 1.0, *ω* = 1.0, − *νω* = 1.0, *ν*(2 − *ω*) = 1.0 (see Methods). Here, new neurites are formed when the neuronal activity exceeds the required level and removed when it falls below it. **(C)** *η* = 5.0, *ψ* = *ϵ* = 15.0, *ν* = 1.0, *ω* = 0.001, −*νω* = 0.001, *ν*(2 − *ω*) = 1.999 (see Methods). Here, the growth curve is shifted up along the y-axis by decreasing the value of *ω*. New neurites are formed when the neuronal activity is less than the homeostatic level and removed (at a very low rate) when it exceeds it.

Here, *ν* is a scaling factor and *η, ϵ* define the width and location of the Gaussian curve on the x-axis. Extending the original MSP framework, we add a new parameter *ω* that controls the location of the curve on the y-axis. The relationship between *η*, *ϵ* and the optimal activity level of a neuron, *ψ*, govern the activity-dependent dynamics of each type of synaptic element. A neuron should not turn over neurites when its activity is optimal ([*Ca*^2+^] = *ψ*). This implies that the growth curves must be placed such that *dz/dt* = 0 when [*Ca*^2+^] = *ψ*. Hence, *ψ* can take one of two values: (*ψ* = *η*) or (*ψ* = *ϵ*), and the turnover of synaptic elements dz/dt is:

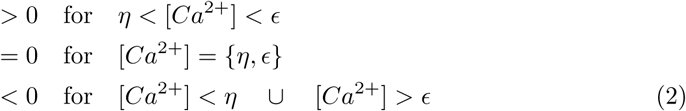

This is illustrated in Fig. 2. Other than in a window between *η* and *ϵ* where new neurites sprout, they retract. The new parameter, *ω*, permits us to adjust the speed of sprouting and retraction (figs. 2B and 2C). In Fig. 2B with (*ψ* = *η*), new neurites will only be formed when the neuron experiences activity that is greater than its homeostatic value (*ψ* < [*Ca*^2+^] < *ϵ*). Fig. 2C, on the other hand, shows the case for (*ψ* = *ϵ*), where growth occurs when neuronal activity is less than optimal (*η* < [*Ca*^2+^] < *ψ*).

The [*Ca*^2+^] for each neuron represents a time averaged measure of its electrical activity:

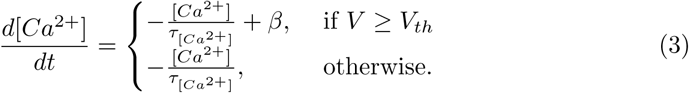

Here, *τ*_[*Ca*^2+^]_ is the time constant with which [*Ca*^2+^] decays in the absence of a spike, *β* is the constant increase in [*Ca*^2+^] caused by each spike, *V* is the membrane potential of the neuron, and *V*_*th*_ is the threshold membrane potential.

Figures 3 and 4 provide an overview of the activity in the network observed in our simulations. The network is initially balanced by the homeostatic inhibitory STDP mechanism, which results in establishing its physiological state where it displays low frequency AI firing similar to cortical neurons [41] (*t* < 1500 s in Figs. 3B, 3C and 4A, and panel 1 in figs. 3A and 4B). Once this AI state is achieved, homeostatic structural plasticity is enabled, and it is confirmed that the network maintains its balanced state under the combined action of the two homeostatic mechanisms (1500 s < *t* < 2000 s in Figs. 3B, 3C and 4A). At (*t* = 2000 s), the network is deafferented by removing external inputs to neurons in the LPZ.

**Fig 3.**
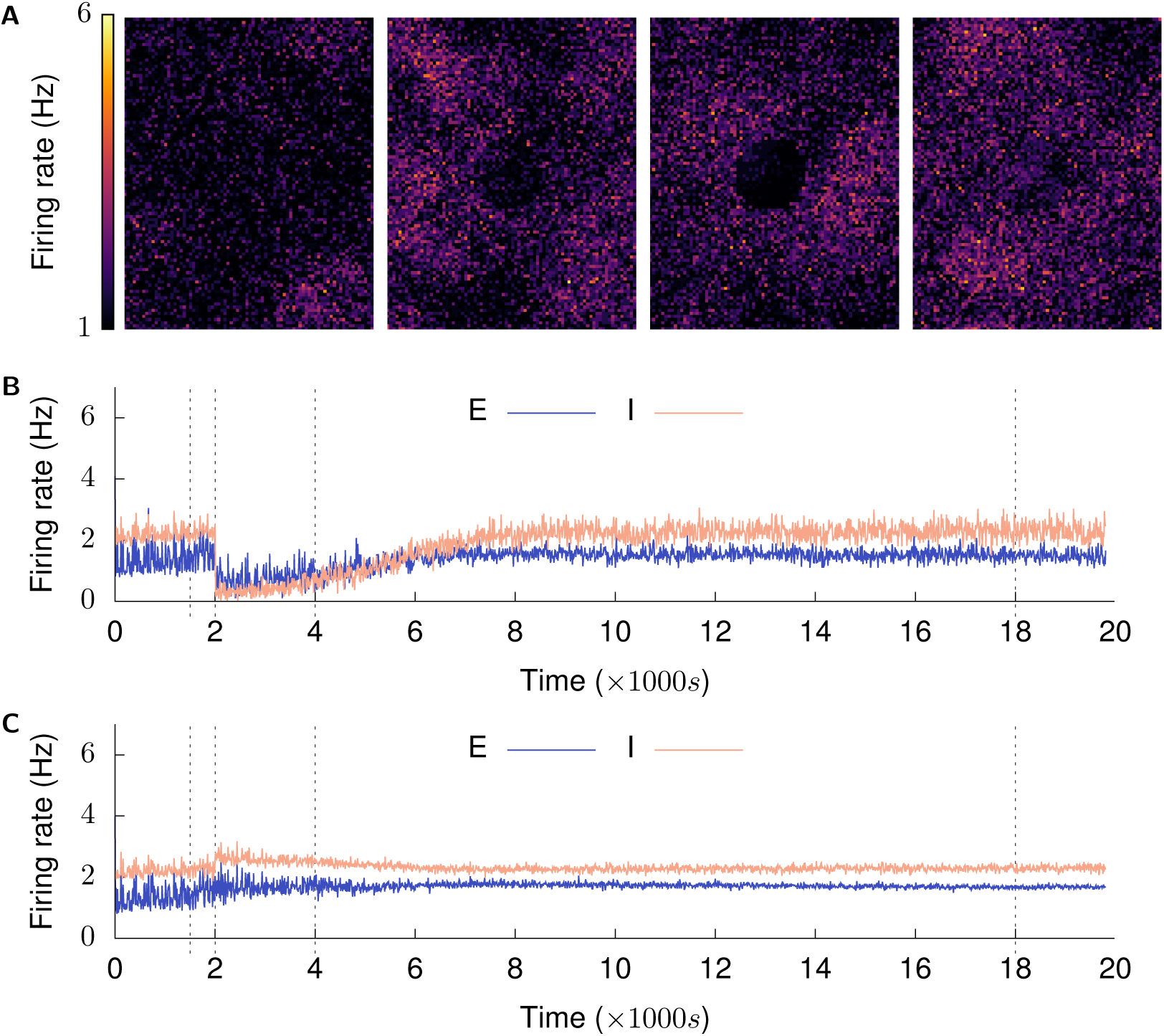
Recovery of activity over time: (Mean firing rates of neurons are calculated over a 2500 ms window): **(A)** shows the firing rates of the whole excitatory population at *t* = {1500 s, 2001.5 s, 4000 s, and 18 000 s}. These are marked by dashed lines in the next graphs. **(B)** shows mean firing rate of neurons in LPZ-C; **(C)** shows mean firing rate of neurons in peri-LPZ; The network is permitted to achieve its balanced Asynchronous Irregular (AI) low frequency firing regime under the action of inhibitory synaptic plasticity (*t* ≤ 1500 s). Our structural plasticity mechanism is then activated to confirm that the network remains in its balanced AI state (panel 1 in Fig. 3A). At (*t* = 2000 s), neurons in the LPZ are deafferented (panel 2 in Fig. 3A is at *t* = 2001.5 s) and the network allowed to repair itself under the action of our structural plasticity mechanism (panels 3 (*t* = 4000 s) and 4 (*t* = 18 000 s) in Fig. 3A).

**Fig 4.**
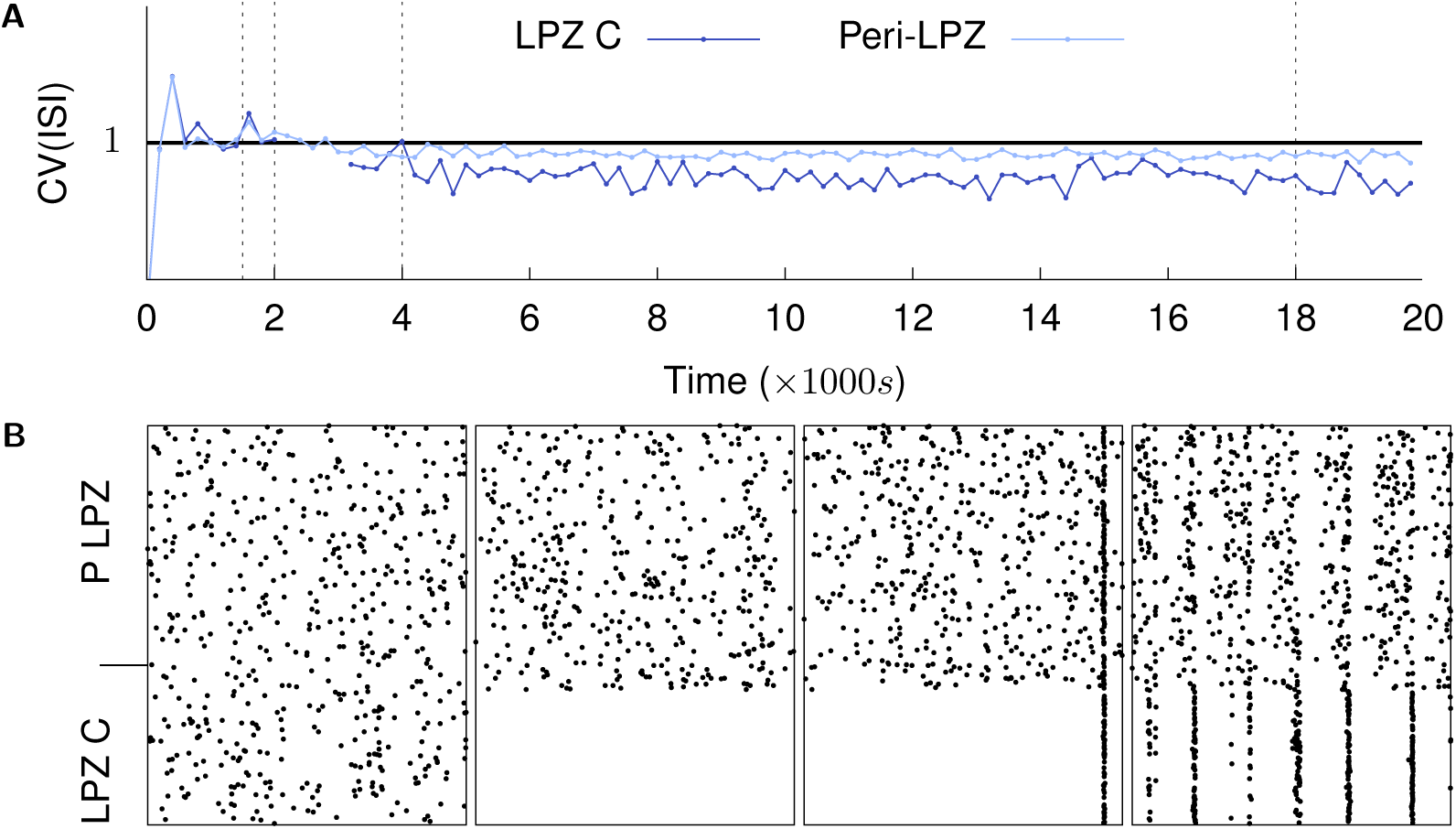
Recovery of activity over time: population firing characteristics: (Mean firing rates of neurons are calculated over a 2500 ms window): **(A)** shows the coefficient of variation (*CV*) of the inter-spike intervals of neurons in the LPZ-C and peri-LPZ. The graph is discontinuous because ISI CV is undefined in the absence of spikes in the LPZ C; **(B)** shows spike times of neurons in the LPZ C and peri-LPZ over a 1 s period at *t* = {1500 s, 2001.5 s, 4000 s, and 18 000 s}. The network is permitted to achieve its balanced Asynchronous Irregular (AI) low frequency firing regime under the action of inhibitory synaptic plasticity (*t* ≤ 1500 s). At (*t* = 2000 s), neurons in the LPZ are deafferented (panel 2 in Fig. 4B is at *t* = 2001.5 s) and the network allowed to repair itself under the action of our structural plasticity mechanism (panels 3 (*t* = 4000 s) and 4 (*t* = 18 000 s) in Fig. 4B). As can be seen here, the network does not return to its AI state after repair.

In line with experimental findings, the immediate result of deafferentation is the loss of activity in neurons of the LPZ. For neurons outside the LPZ, on the other hand, our simulations show an increase in activity suggesting that the net effect of LPZ deafferentation on these neurons is a loss of inhibition rather than excitation (*t* = 2000 s in Fig. 3C). To our knowledge, this phenomenon has not yet been investigated in experiments, and an increase in neuronal activity following deafferentation of a neighbouring area is therefore the first testable prediction provided by our model. The change in activity caused by deafferentation stimulates neurite turnover in neurons of the network in accordance with our proposed activity dependent growth rules (*t* > 2000 s). Over time, activity is gradually restored in the network to pre-deafferentation levels (*t* = 18 000 s in figs. 3B and 3C, and panel 4 in figs. 3A and 4B). In the following sections, we demonstrate that the alterations in network connectivity during repair follow the same regime as reported in experiments, and we derive our growth rules.

Even though the mean activity of neurons within and outside the LPZ returns to pre-deprivation levels, the network reorganization by structural plasticity leads to synchronous spiking in neurons in the LPZ, instead of the AI firing during the pre-deprivation stages in our simulations (*t* > 4000 s in Fig. 4A, and panels 3 and 4 in Fig. 4B). This predicted effect of network rewiring on the temporal characteristics of neural activity should be an interesting subject for future experimental studies. Furthermore, the observed lack of AI activity in the LPZ is expected to have functional implications; this is another promising topic for future theoretical work.

### Activity-dependent dynamics of post-synaptic structures

Since the activity of neurons depends on the inputs received through their post-synaptic neurites, we first derived the growth rules for these neurites. All neurons in the LPZ, excitatory and inhibitory, show near zero activity after deafferentation due to a net loss in excitatory input (panel 2 in figs. 3A and 4B, and *t* = 2000 s in Fig. 3B). Experimental studies report that these neurons gain excitatory synapses on newly formed dendritic spines [28] and lose inhibitory shaft synapses [11] to restore activity after deprivation. The increase in lateral excitatory projections to these neurons requires them to gain excitatory dendritic (postsynaptic) elements to serve as contact points for excitatory axonal collaterals. At the same time, inhibitory synapses can be lost by the retraction of inhibitory dendritic elements. This suggests that new excitatory post-synaptic elements should be formed and inhibitory ones removed when neuronal activity is less than its optimal level (([*Ca*^2+^] < *ψ*) in Fig. 5A):

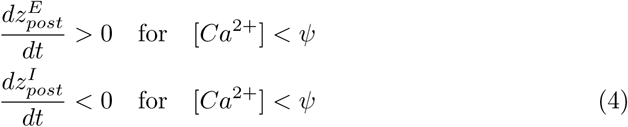

**Fig 5.**
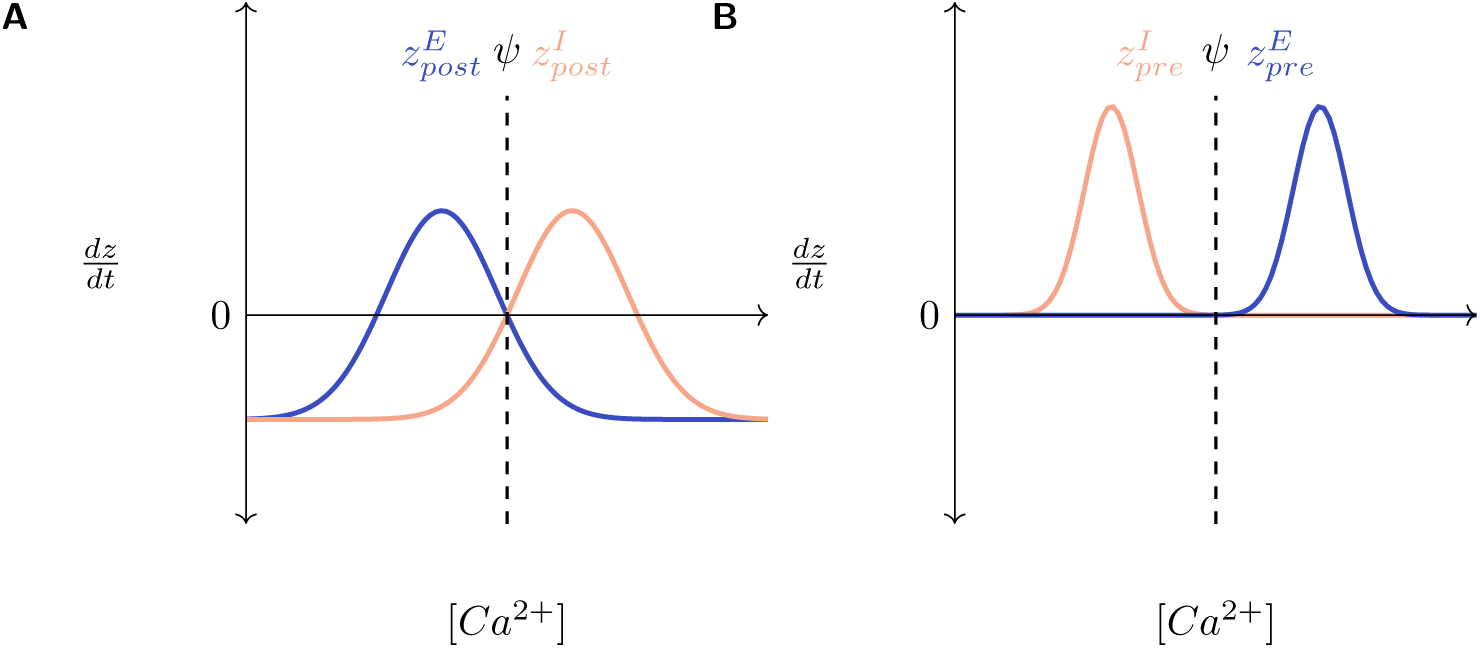
Activity-dependent dynamics of synaptic elements (*dz/dt*) as functions of a neuron’s time averaged activity ([*Ca*^2+^]): **(A)** post-synaptic elements: The balance between excitation and inhibition (E-I balance) received by a neuron may be disturbed by a change in either of the two types of input. Post-synaptic elements of a neuron react to deviations in activity from the optimal level (*ψ*) by countering the changes in excitatory or inhibitory inputs to restore the E-I balance. For both excitatory and inhibitory neurons, excitatory post-synaptic elements sprout when the neuron experiences a reduction in its activity, and retract when the neuron has received extra activity. Inhibitory post-synaptic elements for all neurons follow the opposite rule: they sprout when the neuron has extra activity and retract when the neuron is deprived of activity. **(B)** pre-synaptic elements. In excitatory neurons, axonal sprouting is stimulated by extra activity. In inhibitory neurons, on the other hand, deprivation in activity stimulates axonal sprouting. Synaptic elements that do not find corresponding partners to form synapses (free synaptic elements) decay exponentially with time. These graphs are for illustration only. Please refer to Table 5 for parameter values.

While we were unable to find experimental evidence on the activity of excitatory or inhibitory neurons just outside the LPZ, in our simulations, these neurons exhibit increased activity after deafferentation (*t* = 2000 s in Fig. 3C). Unlike neurons in the LPZ that suffer a net loss of excitation, these neurons appear to suffer a net loss of inhibition, which indicates that they must gain inhibitory and lose excitatory inputs to return to their balanced state. Hence, the formation of new inhibitory dendritic elements and the removal of their excitatory counterparts occurs in a regime where neuronal activity exceeds the required amount (([*Ca*^2+^] > *ψ*) in Fig. 5A):

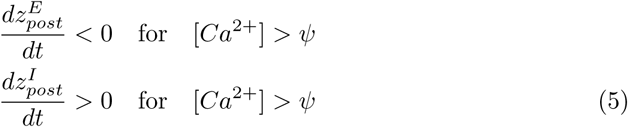

The constraints described by equations Eqs. (2), (4) and (5) can be satisfied by Gaussian growth rules for excitatory and inhibitory dendritic elements, with 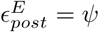 and 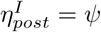, respectively (Fig. 5A and Table 1). Given the distinct characteristics of excitation and inhibition, the two growth rules were treated independently and the parameters governing them were tuned iteratively over multiple simulation runs. For example, sufficiently high values for the rate of formation of inhibitory dendritic elements had to be selected for excitatory neurons to prevent the build up of excessive excitation (Table 5).

**Table 1.** Growth curve parameters for post-synaptic elements.

|  | Excitatory |  |  | Inhibitory |  |  |
| --- | --- | --- | --- | --- | --- | --- |
| | $(\epsilon = \psi)$ | $(\eta < \psi < \epsilon)$ | $(\psi = \eta)$ | $(\epsilon = \psi)$ | $(\eta < \psi < \epsilon)$ | $(\psi = \eta)$ |
| Normal | Yes | No | Yes | Yes | No | Yes |
| Repair | Yes |  | No | No |  | Yes |

Figures 6 and 7 show the time course of rewiring of excitatory and inhibitory connections to excitatory neurons in the centre of the LPZ that results from the growth curves in our simulations. As described in experimental studies, the loss of activity by neurons in the LPZ is followed by an increase in excitatory input connections [13, 28] and a transient reduction in inhibitory input connections [11]. Specifically, as also found in these experiments, the increase in excitatory inputs is dominated by an ingrowth of lateral projections from outside the LPZ. Both of these features can be seen in figs. 6A and 6B. As shown in Figs. 8 and 9, neurons directly outside the LPZ lose excitatory and gain inhibitory input connections to reduce their activity back to their optimal values. Furthermore, in line with experimental observations, a significant contribution to the new inhibitory inputs to these neurons is provided by new inhibitory projections from within the LPZ. Given the small number of inhibitory neurons in the LPZ, however, their inhibitory projections are insufficient to stabilise the large number of neurons outside the LPZ in our simulations. Hence, inhibitory projections are also recruited from inhibitory neurons outside the LPZ.

**Fig 6.**
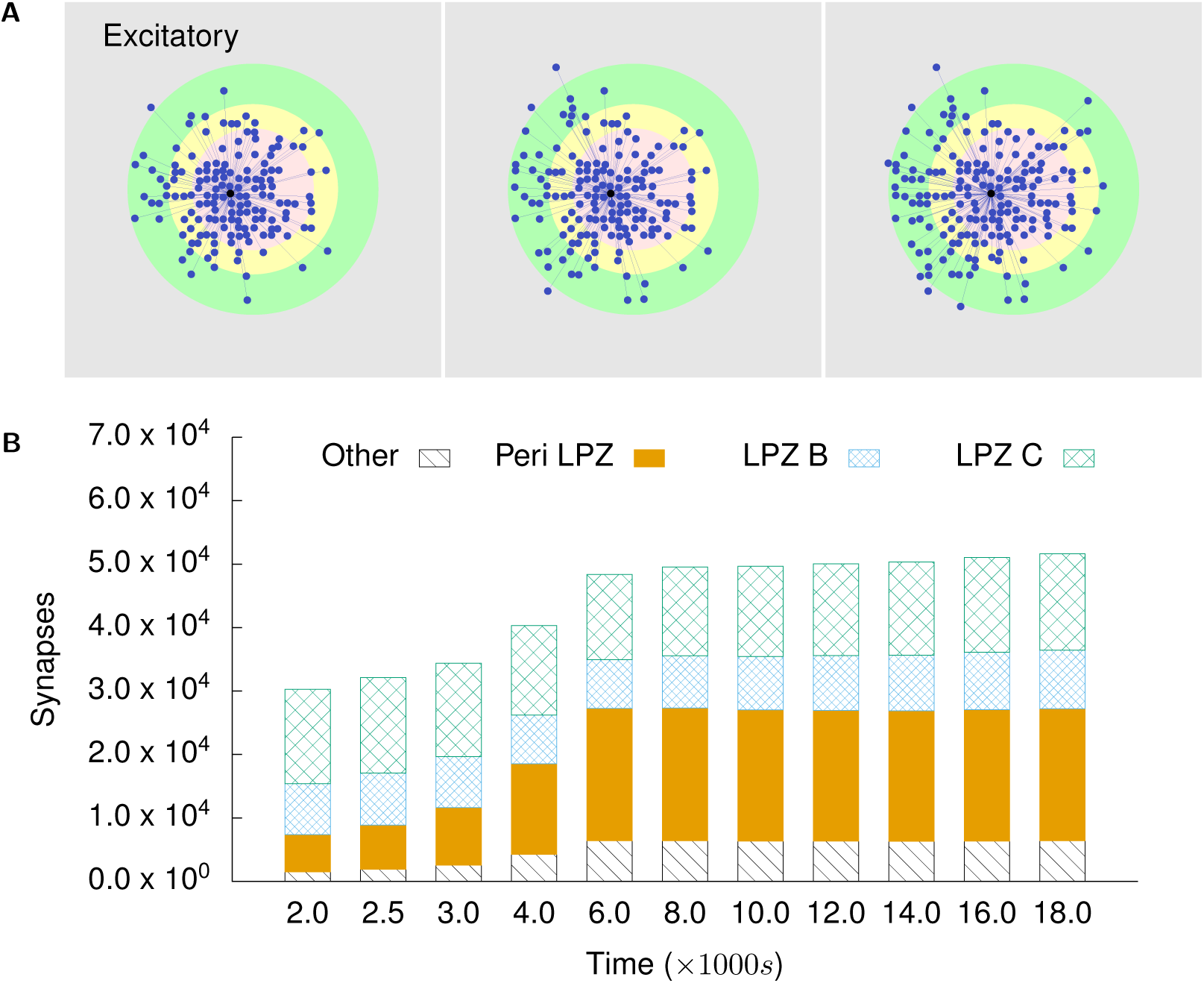
Excitatory projections to excitatory neurons in the centre of the LPZ: **(A)** shows incoming excitatory projections to a randomly chosen neuron in the centre of the LPZ at different stages of our simulations. From left to right: *t* = 2000 s, *t* = 4000 s, and *t* = 18 000 s. **(B)** shows the total numbers of incoming excitatory projections to neurons in the centre of the LPZ from different regions at different points in time. Following our proposed growth rules for post-synaptic elements and consistent with experimental reports, the deprived neurons in the LPZ C gain lateral excitatory inputs from neurons outside the LPZ.

**Fig 7.**
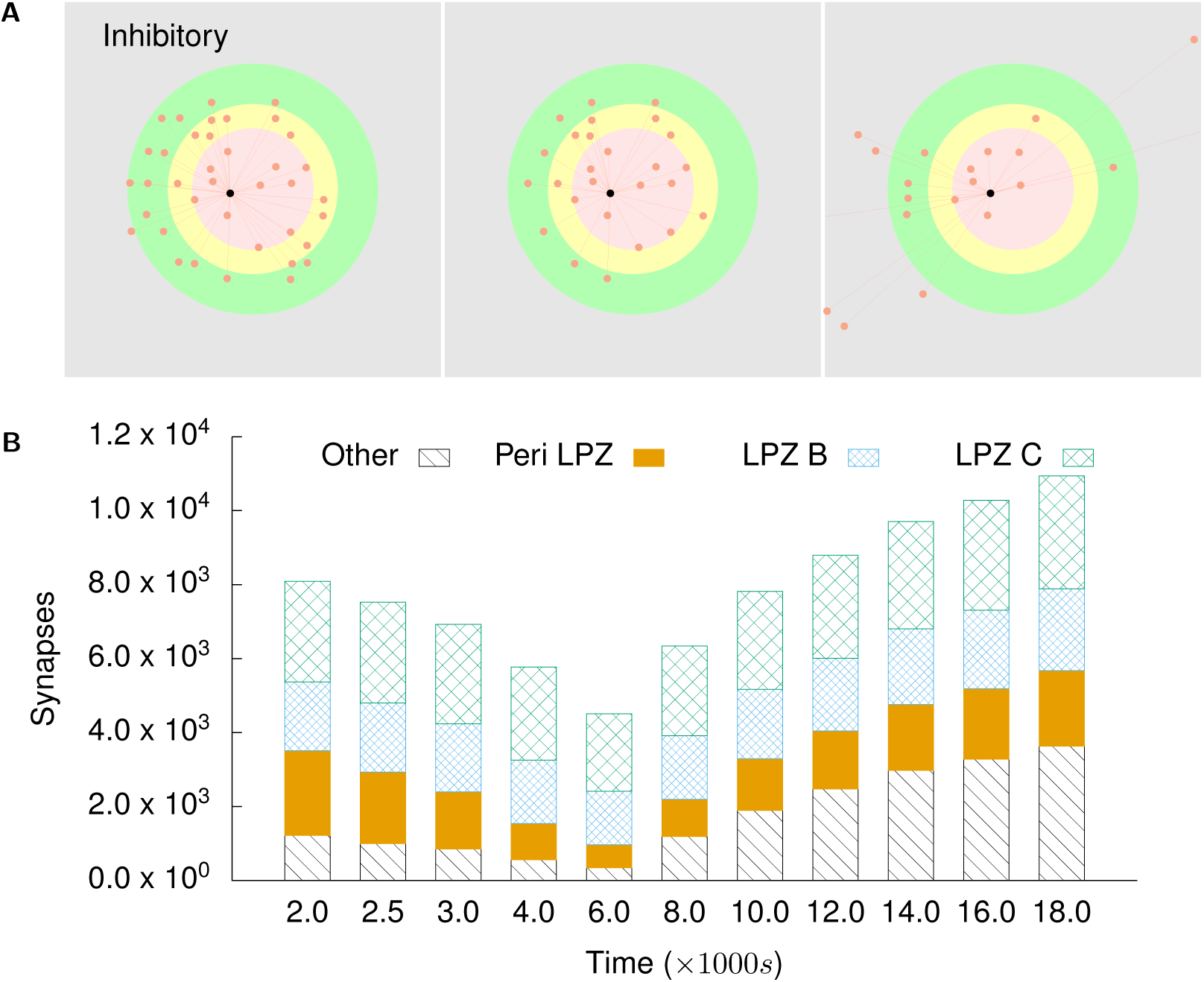
Inhibitory projections to excitatory neurons in the centre of the LPZ: **(A)** shows incoming inhibitory projections to a randomly chosen neuron in the centre of the LPZ at different stages of our simulations. From left to right: *t* = 2000 s, *t* = 4000 s, and *t* = 18 000 s. **(B)** shows the total numbers of incoming inhibitory projections to neurons in the centre of the LPZ from different regions at different points in time. Also in line with biological observations, they temporarily experience disinhibition after deafferentation. However, as these neurons gain activity from their new lateral excitatory inputs, the number of their inhibitory input connections increases again in order to restore the E-I balance.

**Fig 8.**
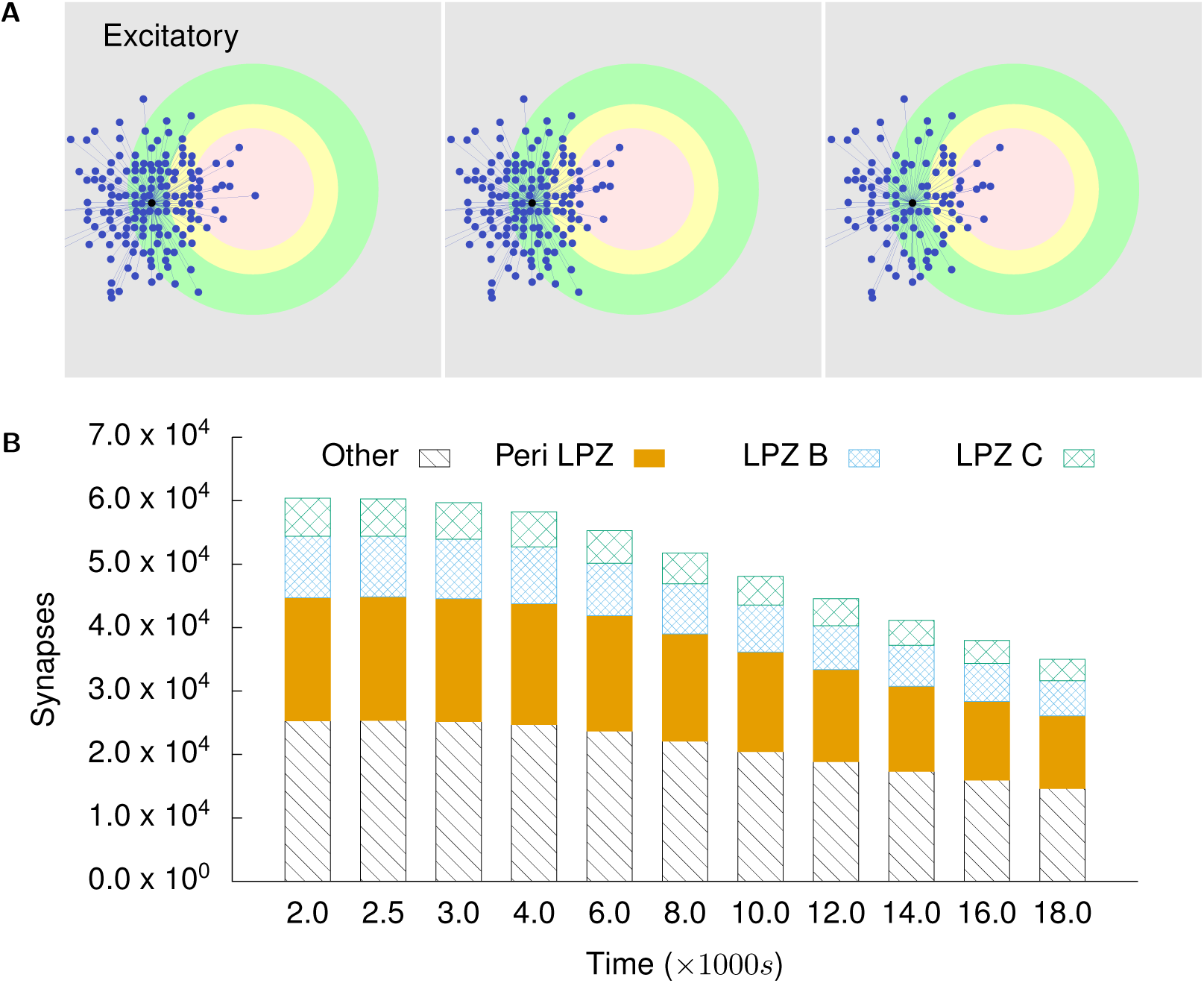
Excitatory projections to excitatory neurons in the peri-LPZ: **(A)** shows incoming excitatory projections to a randomly chosen neuron in the peri-LPZ at different stages of our simulations. From left to right: *t* = 2000 s, *t* = 4000 s, and *t* = 18 000 s. **(B)** shows the total numbers of incoming excitatory projections to neurons in the peri-LPZ from different regions at different points in time. In contrast to neurons in the LPZ, neurons outside the LPZ experience an increase in activity in our simulations. As a result of our growth rules, these neurons lose excitatory inputs.

**Fig 9.**
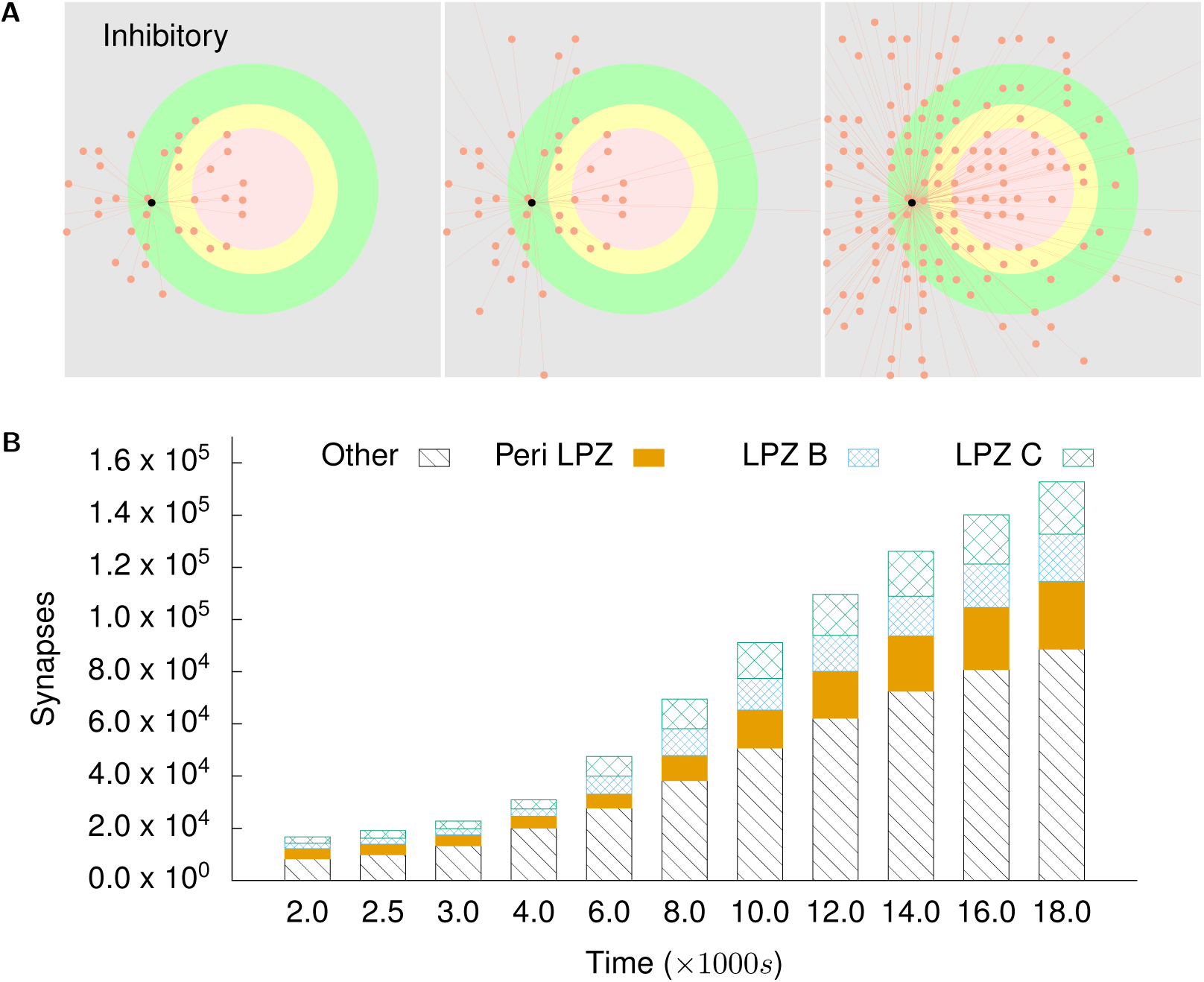
Inhibitory projections to excitatory neurons in the peri-LPZ: **(A)** shows incoming inhibitory projections to a randomly chosen neuron in the peri-LPZ at different stages of our simulations. From left to right: *t* = 2000 s, *t* = 4000 s, and *t* = 18 000 s. **(B)** shows the total numbers of incoming excitatory projections to neurons in the peri-LPZ from different regions at different points in time. In contrast to neurons in the LPZ, neurons outside the LPZ experience an increase in activity in our simulations. As a result of our growth rules, these neurons gain inhibitory inputs.

#### Post-synaptic growth rules stabilise individual neurons

Experimental evidence suggests that not just networks, but also individual neurons in the brain maintain a finely tuned balance between excitation and inhibition (E-I balance) [44–46]. This raises the question whether the complementary nature of our excitatory and inhibitory post-synaptic growth rules is sufficient to ensure stability at the level of single neurons.

Since the state of each neuron is tightly coupled to the states of other neurons in the network, we modelled a neuron in isolation to investigate how its input connectivity would be affected by changes in activity as per our post-synaptic growth curves (Fig. 10A). The neuron is initialised with an input connectivity similar to a neuron from the network in its steady state: it has the same number of excitatory 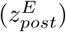 and 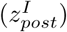 dendritic elements and receives the same mean conductances through them (*g*_*EE*_, *g*_*IE*_). Thus, the [*Ca*^2+^] of the neuron in this state represents its optimal activity (*ψ* = [*Ca*^2+^] at *t* = 0 s in Fig. 10B). In this scenario, the net input conductance received by the neuron (*g*_*net*_), which modulates its activity, can be estimated as the difference of the total excitatory (*g*_*E*_) and inhibitory (*g*_*I*_) input conductances.

**Fig 10.**
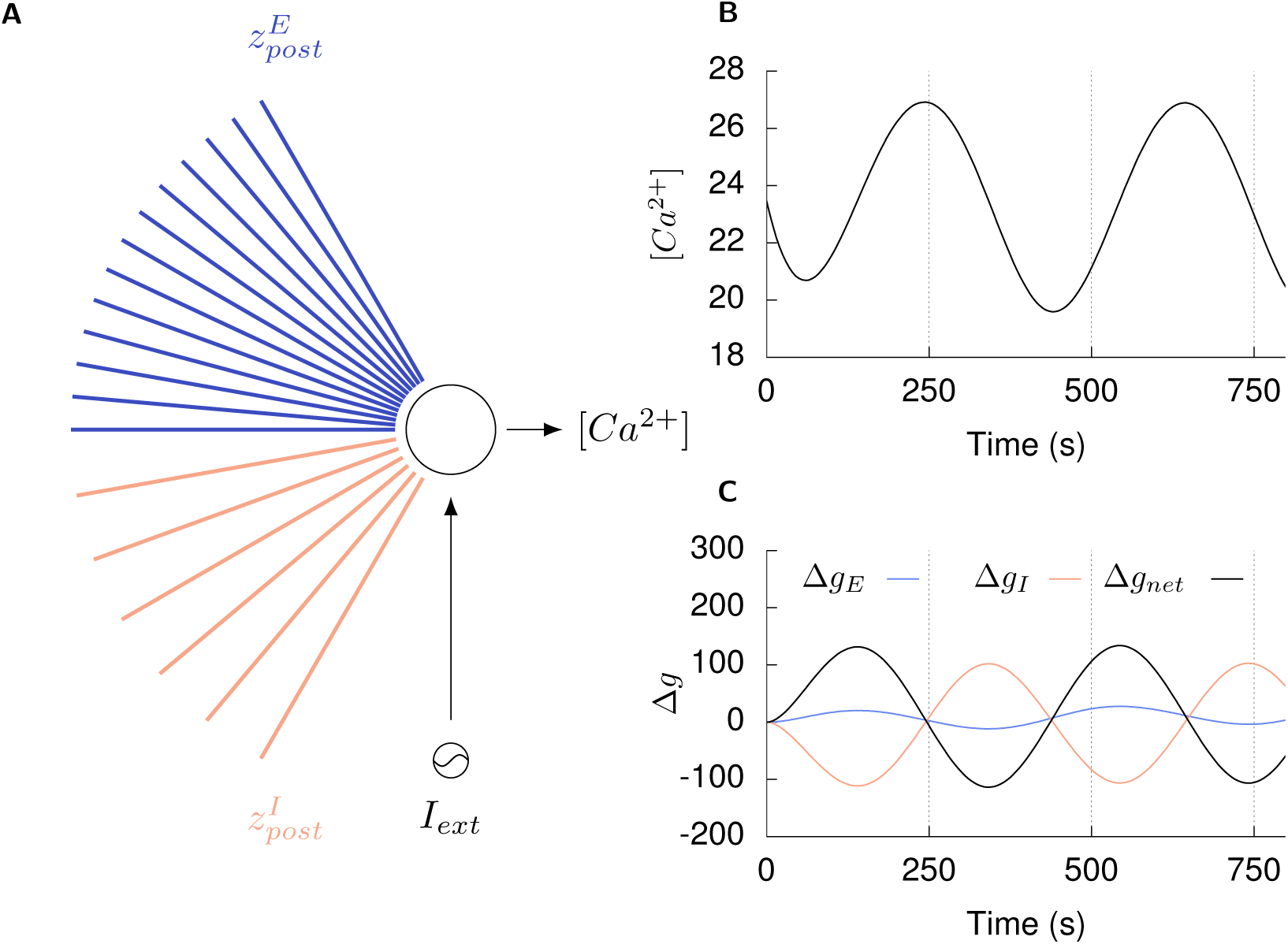
Single neuron simulations show the homeostatic effect of the post-synaptic growth rules: **(A)** A neuron in its steady state receives excitatory (*g*_*E*_) and inhibitory (*g*_*I*_) conductance inputs through its excitatory 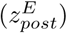 and inhibitory 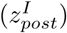 dendritic elements, respectively, such that its activity ([*Ca*^2+^]) is maintained at its optimal level (*ψ*) by its net input conductance (*g*_*net*_). **(B)** An external sinusoidal current stimulus (*I*_*ext*_) is applied to the neuron to vary its activity from the optimal level. **(C)** Under the action of our post-synaptic growth curves, the neuron modifies its dendritic elements to change its excitatory (∆*g*_*E*_) and inhibitory (∆*g*_*I*_) conductance inputs such that the net change in its input conductance (∆*g*_*net*_) counteracts the change in its activity: an increase in [*Ca*^2+^] due to the external stimulus is followed by a decreas in net input conductance through the post-synaptic elements and vice versa (dashed lines in Figs. 10B and 10C).

The activity of the neuron is then varied by an external sinusoidal current stimulus (Fig. 10B). In addition, the deviation of the neuron’s excitatory (∆*g*_*E*_), inhibitory (∆*g*_*I*_), and net input conductance (∆*g*_*net*_) from baseline levels due to the formation or removal of dendritic elements under the action of the growth curves is recorded (Fig. 10C). We find that that modifications of the input connectivity of the neuron result in alterations to its excitatory and inhibitory input such that the net change in its input conductance counteracts changes in its activity: an increase in [*Ca*^2+^] due to the external stimulus is followed by a decrease in net input conductance through the post-synaptic elements and vice versa (dashed lines in figs. 10B and 10C). These simulation results show that even though the activity dependent growth rules of excitatory and inhibitory post-synaptic elements are derived from network simulations, they also serve a homeostatic function in single neurons.

### Activity dependent dynamics of pre-synaptic structures

While the activity dependent formation and degradation of post-synaptic elements provides a homeostatic mechanism for the stabilisation of activity in single neurons and the network, the increase in excitatory or inhibitory input received by a neuron also relies on the availability of pre-synaptic counterparts. We derive activity dependent growth rules for excitatory 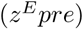 and inhibitory 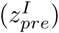 pre-synaptic elements in a similar manner to that used for post-synaptic elements.

Within the LPZ, the increase in excitation requires a corresponding increase in the supply of excitatory pre-synaptic elements. Experimental evidence reports a sizeable increase in the formation and removal of axonal structures in and around the LPZ [27], with a marked addition of lateral projections from neurons outside the LPZ into it [6]. While an increase in pre-synaptic elements within the LPZ may contribute to repair, an inflow of activity from the periphery of the LPZ to its centre has been observed in experiments [6, 20, 28], pointing to the inwards sprouting of excitatory axonal projections from outside the LPZ as the major driver of homeostatic rewiring. For this sprouting of excitatory projections from the non-deafferentated area into the LPZ to take place in our simulations, the increase in activity in neurons outside the LPZ must stimulate the formation of their excitatory axonal elements:

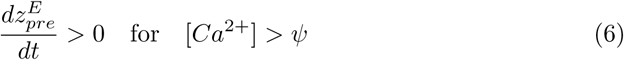

Conversely, neurons outside the LPZ with increased activity need access to inhibitory pre-synaptic elements in order to receive the required additional inhibitory input. Deafferentation studies in mouse somatosensory cortex [6] report more than a 2.5 fold increase in the lengths of inhibitory axons projecting out from inhibitory neurons in the LPZ two days after the peripheral lesion. This outgrowth of inhibitory projections preceded and was faster than the ingrowth of their excitatory analogues [6, 9]. In our simulations, the experimentally observed outward protrusion of inhibitory axons from the LPZ requires that the formation of inhibitory pre-synaptic elements is driven by reduced neuronal activity:

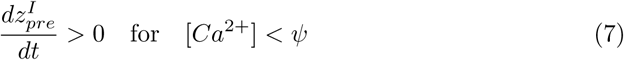

To further validate the derived pre-synaptic growth curves, shown in Fig. 5B and Table 2, the complete set of possible pre-synaptic growth curves was tested. These are labelled **G0, G1, G2, G3, G4**, and **G5** and illustrated in Fig. 11:

**Table 2.** Growth curve parameters for pre-synaptic elements.

|  | Excitatory |  |  | Inhibitory |  |  |
| --- | --- | --- | --- | --- | --- | --- |
| | $(\epsilon = \psi)$ | $(\eta < \psi < \epsilon)$ | $(\psi = \eta)$ | $(\epsilon = \psi)$ | $(\eta < \psi < \epsilon)$ | $(\psi = \eta)$ |
| Normal | Yes | No | Yes | Yes | No | Yes |
| Repair | No |  | Yes | Yes |  | No |

**Fig 11.**
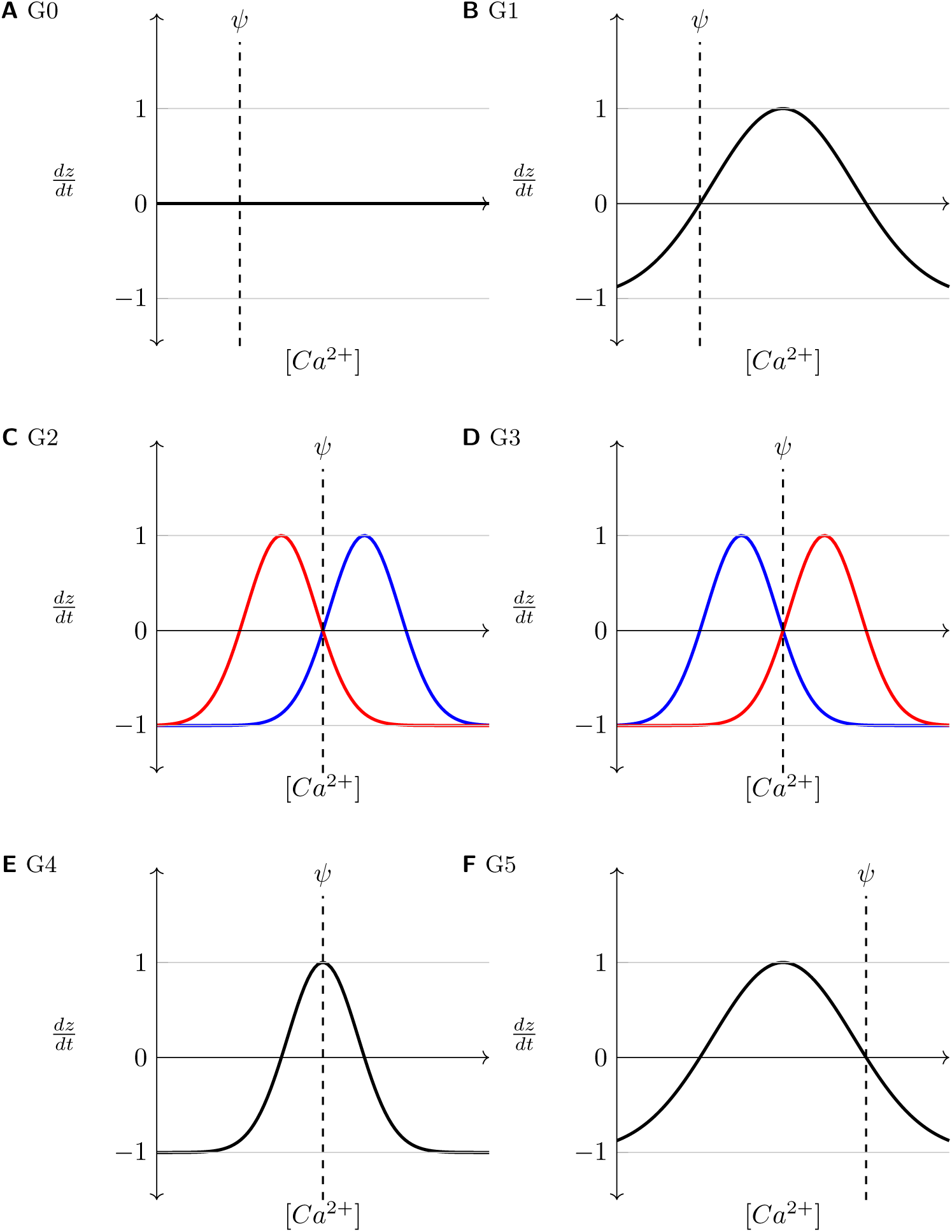
Axonal growth curves investigated in the study. (where applicable, Red: inhibitory, Blue: excitatory).

- **G0**: control case where there are no growth curves, achieved by setting *ν* = 0,
- **G1**: both inhibitory and excitatory axons sprout when activity is more than required,
- **G2**: (the selected growth curves shown in Fig. 5B fall into this category) inhibitory axons sprout when activity is less than optimal, but excitatory axons sprout when activity is more than required,
- **G3**: excitatory axons sprout when activity is less than optimal, but inhibitory axons sprout when activity is more than required,
- **G4**: both excitatory and inhibitory axons sprout at optimal activity, and
- **G5**: both inhibitory and excitatory axons sprout when activity is less than optimal.

As summarised in Table 3, only the previously derived pre-synaptic growth curves reproduced all experimentally reported features of the repair process. While a few other pre-synaptic growth curves did allow simulations to show an increase in activity in the LPZ and a loss of activity outside it, the networks in these simulations did not re-balance to a stable state.

**Table 3.** Summary of axonal growth curves tested in the model.

|  | G0 | G1 | G2 | G3 | G4 | G5 |
| --- | --- | --- | --- | --- | --- | --- |
| Initially remains stable | Y | Y | Y | Y | N | Y |
| LPZ gains activity | Y | Y | Y | N | NA | N |
| Outside LPZ loses activity | Y | Y | Y | NA | NA | NA |
| Returns to balanced state | N | N | Y | NA | NA | NA |
| LPZ B restores before LPZ C | NA | NA | Y | NA | NA | NA |
| Ingrowth of excitatory projections | NA | NA | Y | NA | NA | NA |
| Outgrowth of inhibitory projections | NA | NA | Y | NA | NA | NA |
| Disinhibition in LPZ | NA | NA | Y | NA | NA | NA |

Similar to the post-synaptic growth rules, the pre-synaptic growth rules for excitatory and inhibitory neurons were also treated separately and their parameters were tuned iteratively over repeated simulations. Since inhibitory neurons form only one-fourth of the neuronal population, and only a small number of these fall into the LPZ, in this study, simulations require the growth rates of inhibitory axonal elements to be high enough to stabilise the large number of hyperactive neurons outside the LPZ (Table 5).

Figures 12A and 12B show the rewiring of axonal projections from an excitatory neuron in the peri-LPZ and an inhibitory neuron in the centre of the LPZ, respectively. Following the growth functions derived above, our simulations correctly reproduce the inward sprouting of excitatory axons into the LPZ and the outward sprouting of inhibitory axons from the LPZ that is observed during the repair process.

**Fig 12.**
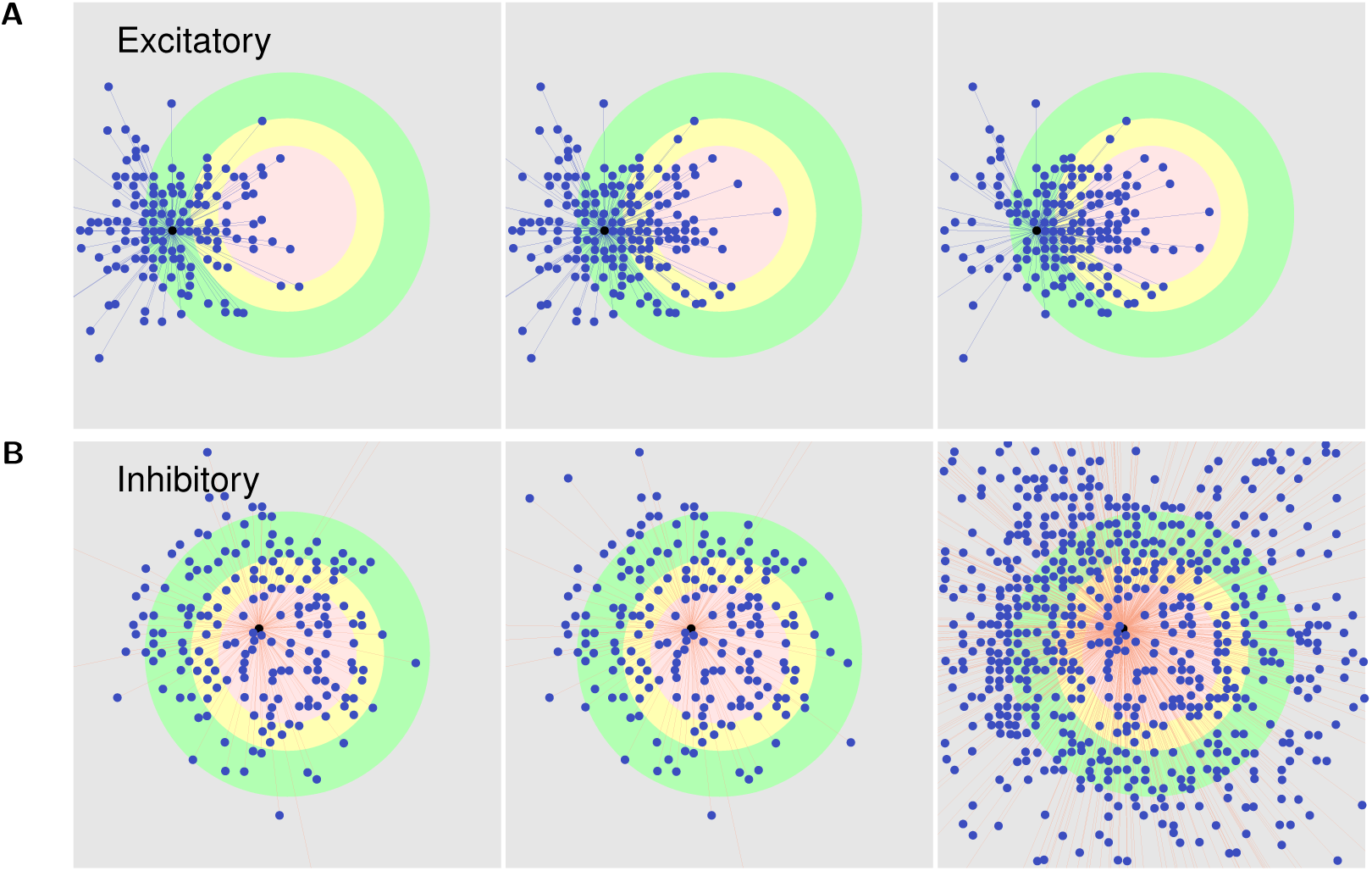
Outgoing projections: **(A)** shows the outgoing (axonal) projections of an excitatory neuron in the peri-LPZ. **(B)** shows the outgoing (axonal) projections of an inhibitory neuron in the LPZ C. From left to right: *t* = 2000 s, *t* = 4000 s, and *t* = 18 000 s. As per our suggested growth rules for pre-synaptic elements, excitatory neurons produce new pre-synaptic elements and sprout axonal projections when they experience extra activity, while inhibitory neurons form new pre-synaptic elements and grow axons when they are deprived of activity. As a consequence and in line with experimental data, following deafferentation of the LPZ, excitatory neurons in the peri-LPZ sprout new outgoing projections that help transfer excitatory activity to neurons in the LPZ. Also in accordance with experimental work, inhibitory neurons inside the LPZ form new outgoing connections that transmit inhibition to neurons outside the LPZ.

### Synaptic and structural plasticity are both necessary for repair

In all our previous simulations, the network rewiring after deafferentation of the LPZ occurred in the presence of both activity-dependent structural plasticity and inhibitory synaptic plasticity. These results show that both types of homeostatic plasticity can co-exist during successful network repair, but they do not indicate their respective contributions to restoring activity in the network. In order to study the functional role of the two plasticity mechanisms in the homeostatic regulation of activity after peripheral lesions, we simulated our model with each the mechanisms enabled in isolation (see Methods).

Results from our simulations where structural plasticity is disabled suggest that inhibitory synaptic plasticity alone, while able to re-balance neurons outside the LPZ by increasing the strength of their inhibitiory inputs, fails to restore activity in the deprived neurons in the LPZ even after small peripheral lesions (figs. 13A and 13D). Although the homeostatic inhibitory synaptic plasticity on its own leads to a reduction in conductances of the inhibitory synapses projecting onto neurons in the LPZ, this is not sufficient to reactivate them. The stabilisation of activity in the neurons outside the LPZ, however, is successful due to the strengthening of IE synapses by STDP. In the absence of network rewiring by structural plasticity, this leads to a network where the neurons outside the LPZ retain their functionality while the LPZ is effectively lost. This indicates that the larger deviations from the desired activity that result from deafferentation in our balanced network model require the reconfiguration of network connectivity by structural plasticity to re-establish a functional balance.

**Fig 13.**
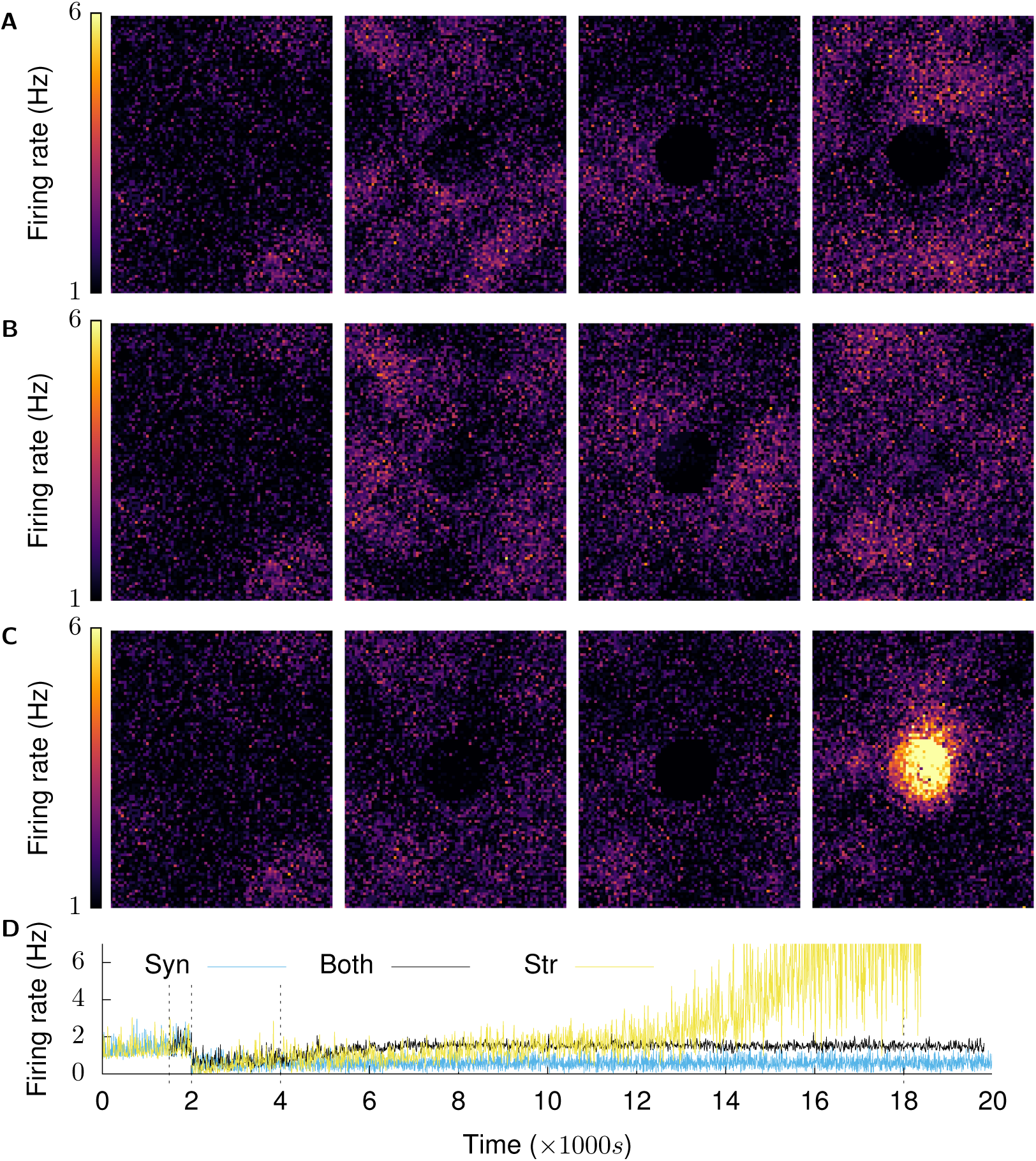
Both structural and synaptic plasticity are required for restoration of activity after deafferentation: **(A), (B), (C)** show firing rate snapshots of neurons at *t* = 1500 s, 2001.5 s, 4000 s, 18 000 s. **(A)** Synaptic plasticity only: after the network has settled in its physiological state by means of synaptic plasticity, structural plasticity is not enabled. With only synaptic plasticity present, the network is unable to restore activity to neurons in the LPZ. Neurons outside the LPZ return to their balanced state, but the neurons in the LPZ are effectively lost to the network. **(B)** Both structural and synaptic plasticity are enabled: neurons in the LPZ regain their low firing rate as before deafferentation. **(C)** Structural plasticity only: after the network has settled in its physiological state by means of synaptic plasticity, homeostatic synaptic plasticity is turned off and only structural plasticity is enabled. With only structural plasticity present, activity returns to neurons in the LPZ but does not stabilise in a low firing rate regime. The firing rate of these neurons continues to increase and, as a result, these neurons continue to turn over synaptic elements. This cascades into increased activity in neurons outside the LPZ, further causing undesired changes in network connectivity. **(D)** shows the mean population firing rates of neurons in the centre of the LPZ for the three simulation configurations. (Panel 1 is identical in all three simulation configurations because the same parameters are used to initialise all simulations.)

Simulations where homeostatic synaptic plasticity was disabled, on the other hand, also failed to re-establish the balanced state of the network before the peripheral lesion (figs. 13C and 13D). While the activity of the deprived neurons in the LPZ initially increased back to pre-lesion levels, under the action of structural plasticity only, the network eventually started exhibiting abnormally high firing rates instead of settling in the desired low firing rate regime. These results suggest that inhibitory synaptic plasticity is required to finely tune inputs to neurons so that the network can achieve its balanced state.

Thus, our simulations predict that both homeostatic processes are required for successful repair—structural plasticity for larger changes in network connectivity and synaptic plasticity for the fine tuning of conductances that establishes stable activity in the network. These results support the idea that multiple plasticity mechanisms work in harmony to sustain functional brain networks at varying time scales.

## Discussion

A better understanding of the factors that influence dynamic alterations in the morphology and connectivity of neuronal axons and dendrites is necessary to improve our knowledge of the processes that shape the development and reorganisation of neuronal circuitry in the adult brain. Here, we present a new, spiking neural network model of peripheral lesioning in a simplified cortical balanced asynchronous irregular network (Figs. 1 and 2). We show that our simulations reproduce the time course of changes in network connectivity as reported in experimental work (Fig. 3), and we provide a number of testable predictions.

First, our model suggests that deafferentation does not necessarily result in the loss or even a decrease of activity in all neurons of the network. Neurons outside the LPZ experience a gain in activity because of a net loss in inhibition in our simulations. This prediction should be tested in future experiments that investigate neuronal activity just outside the LPZ.

Secondly, our model suggests that while the network may restore its mean activity, the temporal fine structure of the activity, and in particular the AI firing characteristic of the network are permanently disturbed by deafferentation. This change in firing patterns of the network also merits experimental validation, especially given its implications for network function. Synchronous firing in the network may not be evident in studies of the mapping between peripheral inputs and network activity. However, in combination with the change in network connectivity, it can affect other types of network function, such as the storage and recall of associative memory. By storing Hebbian assemblies in the network and testing their recall after deafferentation and repair, we are currently exploring this phenomenon.

Thirdly, as the main objective of our work, we suggest different growth rules for differnt types of neurite (Fig. 5). While derived from network lesion experiments that were not aimed at studying the relation between activity and neurite turnover [6, 9, 10, 13, 27–30], evidence from other work seems to support our proposals. Our growth rule for excitatory dendritic elements is coherent with results from an experimental study in hippocampal slice cultures. In their study, Richards et al. note that reduced neuronal activity resulted in the extension of glutamate receptor-dependent processes from dendritic spines of CA1 pyramidal neurons [47]. Furthermore, the predicted growth function for inhibitory dendritic elements is supported by a study by Knott et. al [3], which reports an increase in inhibitory inputs to spines in adult mice after their activity was increased by whisker stimulation [3].

On the pre-synaptic side, axonal turnover and guidance has been investigated in much detail, and is known to be a highly complex process incorporating multiple biochemical pathways [48, 49]. Our hypothesis regarding excitatory pre-synaptic structures is supported by a report by Perez et al. who find that CA1 pyramidal cells, which become hyper-excitable following hippocampal kainate lesions, sprout excitatory axons that may contribute to the epileptiform activity in the region [50]. For inhibitory pre-synaptic elements, we refer to Schuemann et al. who report that enhanced network activity reduced the number of persistent inhibitory boutons [51] over short periods of time (30 minutes) in organotypic hippocampal slice cultures. However, these experiments also found that prolonged blockade of activity (over seven days) did not affect inhibitory synapses, contrary to the reports from peripheral lesion studies [11, 30].

Indirect evidence on the temporal evolution of inhibitory projections to neurons in the LPZ further supports the inhibitory growth rules in our model (Fig. 7B). While an initial disinhibition aids recovery in these deprived neurons, as activity is restored, a subsequent increase in inhibition in our simulations re-establishes the E-I balance in the deafferented region. This is in line with evidence that the pharmacological reduction of inhibition restores structural plasticity in the visual cortex [52]. Our simulations, therefore, support the proposed role of inhibition as control mechanism for the critical window for structural plasticity [15, 53–57].

Our simulation results do not imply that these are the only activity dependent growth rules that can underlie the turnover of neurites. Given the variety of neurons in the brain, many families of growth rules may apply to neurons. For example, Butz and van Ooyen proposed a different set of growth rules using a model of peripheral lesioning in fast spiking neurons that did not investigate the low firing AI state [33]. Different growth rules could therefore apply to brain regions with different neuronal types and firing characteristics.

Finally, our simulation results indicate that the suggested growth rules, while derived from network simulations, can contribute to the stability of activity in individual neurons (Fig. 10). Since structural plasticity and synaptic plasticity are not independent processes in the brain, this is not a wholly surprising result. Structural plasticity of the volumes of spines and boutons underlies the modulation of synaptic efficacy by synaptic plasticity. Thus, given that synaptic plasticity mechanisms can stabilise the firing of individual neurons [58, 59], it follows that structural plasticity mechanisms could also be involved. Further, extending from the functional coupling of synaptic and structural plasticity, our simulations also require both structural and synaptic plasticity for successful network repair (Fig. 13). Thus, our simulation results lend further support to the notion that multiple plasticity mechanisms function in a cooperative manner in the brain.

As a computational modelling study, our work necessarily suffers from various limitations. For example, while the use of simple conductance based point neurons [43] is sufficient for our network study, perhaps even necessary for its tractability [60], it also limits our work. Unlike in the brain where calcium is compartmentalised in neurons [61], a single compartment point neuron model only allows one value of [*Ca*^2+^] for all neurites in a neuron. Thus, each of the neurons in our model can only either sprout or retract a type of neurite at a point in time. This is not the case in biology where different parts of the neuron can undergo structural changes independently of each other. The growth regimes suggested in our work must be understood to address the net formation or removal of neurites only. Furthermore, since a simultaneous homeostatic regulation of different neuronal compartments would be expected to have a larger stabilising effect on the overall activity of the neuron, a single compartment neuron model may also limit the homeostatic effect of the structural plasticity mechanism. Point neurons also lack morphology, and our model is therefore unable to explicitly include the directional formation or removal of synapses. Axonal and dendritic arbors are not explicitly modelled and the directional turnover of synapses that represents axonal sprouting emerges merely from the numbers of connecting partner neurites. Additionally, while it was enough for neurons in our model to be distributed in a two dimensional grid to include a spatial component, this is clearly not true for the brain. Thus, while our model provides a simplified high level view, the investigation of our proposed activity dependent growth rules in more detailed models is an important avenue for future research.

Finally, this work, and computational modelling of structural plasticity in general, are limited by the lack of supporting simulation tools. Most current simulators are designed for network modelling where synaptic connectivity remains constant. Even the NEST simulator [62], where the internal data structures are sufficiently flexible to allow for modification of synapses during simulation [63], currently includes a limited implementation of the MSP algorithm [38]. To incorporate the missing pieces— spatial information and different network connectivity modification strategies, for example—we were required to repeatedly pause simulations to make connectivity updates. This is far less efficient than NEST handling these changes in connectivity internally during continuous simulation runs and added a large overhead to the computational costs of our simulations. The development of companion tools for modelling structural plasticity is however, gradually gaining traction [64] with discussions to allow NEST to communicate with stand alone structural plasticity tools via interfaces such as Connection Set Algebra [65] ongoing.

In conclusion, we present a new general model of peripheral lesioning and repair in simplified cortical spiking networks with biologically realistic AI activity that provides several experimentally testable predictions.

## Methods

We build on and extend the MSP [33] framework to model the activity dependent dynamics of synaptic elements. We developed our new model using the NEST neural simulator [66, 67]. NEST includes an early, partial implementation of the MSP [38]. It does not, for example, currently take spatial information into account while making connectivity updates. More importantly, at this time, the design of the C++ codebase also does not provide access to the lower level rules governing updates in connectivity via the Python API. Making modifications to these to execute new structural plasticity connectivity rules, therefore, requires non-trivial changes to the NEST kernel. Given that work is on-going to modularise the implementation of structural plasticity in NEST such that the computation of changes in connectivity will be left to stand-alone tools that will communicate them to the simulator using interfaces such as the Connection Set Algebra [65] (private communications with the NEST development team), we resorted to disabling connectivity updates in NEST. Instead, we generate connectivity based on our new hypotheses using native Python methods, and use the methods available in PyNEST to modify them in simulations. Our modified version of the NEST source code is available in our fork of the simulator available in a public repository here.

To honour our commitment to Open Science [68], we only made use of Free/Open source software for our work. The complete source code of all simulations run in this work are available on GitHub here. The scripts used to analyse the data generated by the simulation are available in a separate GitHub repository here. These repositories are licensed under the Gnu GPL license (version 3 or later). The data generated by the simulations used in this paper, along with the individual GNUPlot scripts used to generate each figure, will be made available on Zenodo.

### Neuron model

Neurons are modelled as leaky integrate and fire conductance based point neurons with exponential conductances [43], the membrane potentials of which are governed by:

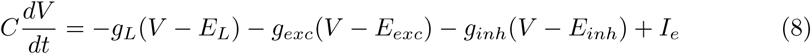

where *C* is the membrane capacitance, *V* is the membrane potential, *g*_*L*_ is the leak conductance, *g*_*exc*_ is the excitatory conductance, *g*_*inh*_ is the inhibitory conductance, *E*_*L*_ is the leak reversal potential, *E*_*exc*_ is the excitatory reversal potential, *E*_*inh*_ is the inhibitory reversal potential, and *I*_*e*_ is an external input current. Incoming spikes induce a post-synaptic change of conductance that is modelled by an exponential waveform following the equation:

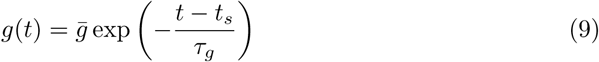

where *τ*_*g*_ is the decay time constant and 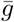 is the maximum conductance as the result of a spike at time *t*_*s*_. Table 4 enumerates the constants related to the neuron model.

Each neuron possesses sets of both pre- and post-synaptic synaptic elements, the total numbers of which are represented by (*z*_*pre*_) and (*z*_*post*_) respectively. Excitatory and inhibitory neurons only possess excitatory 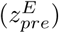 and inhibitory axonal elements 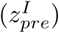 respectively, but they can each host both excitatory and inhibitory dendritic elements (*z*_*post,E*_, *z*_*post,I*_) (since the number of neurites must be a non-negative integer, the floor value of the continuous variable is used for connectivity updates). As in MSP, we model the rate of change of each type of synaptic element, (*dz/dt*), as a Gaussian function of the neuron’s “Calcium concentration” ([*Ca*^2+^]):

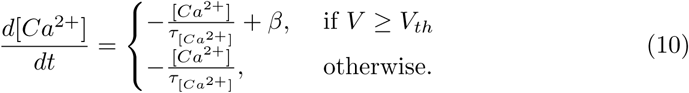

**Table 4.**
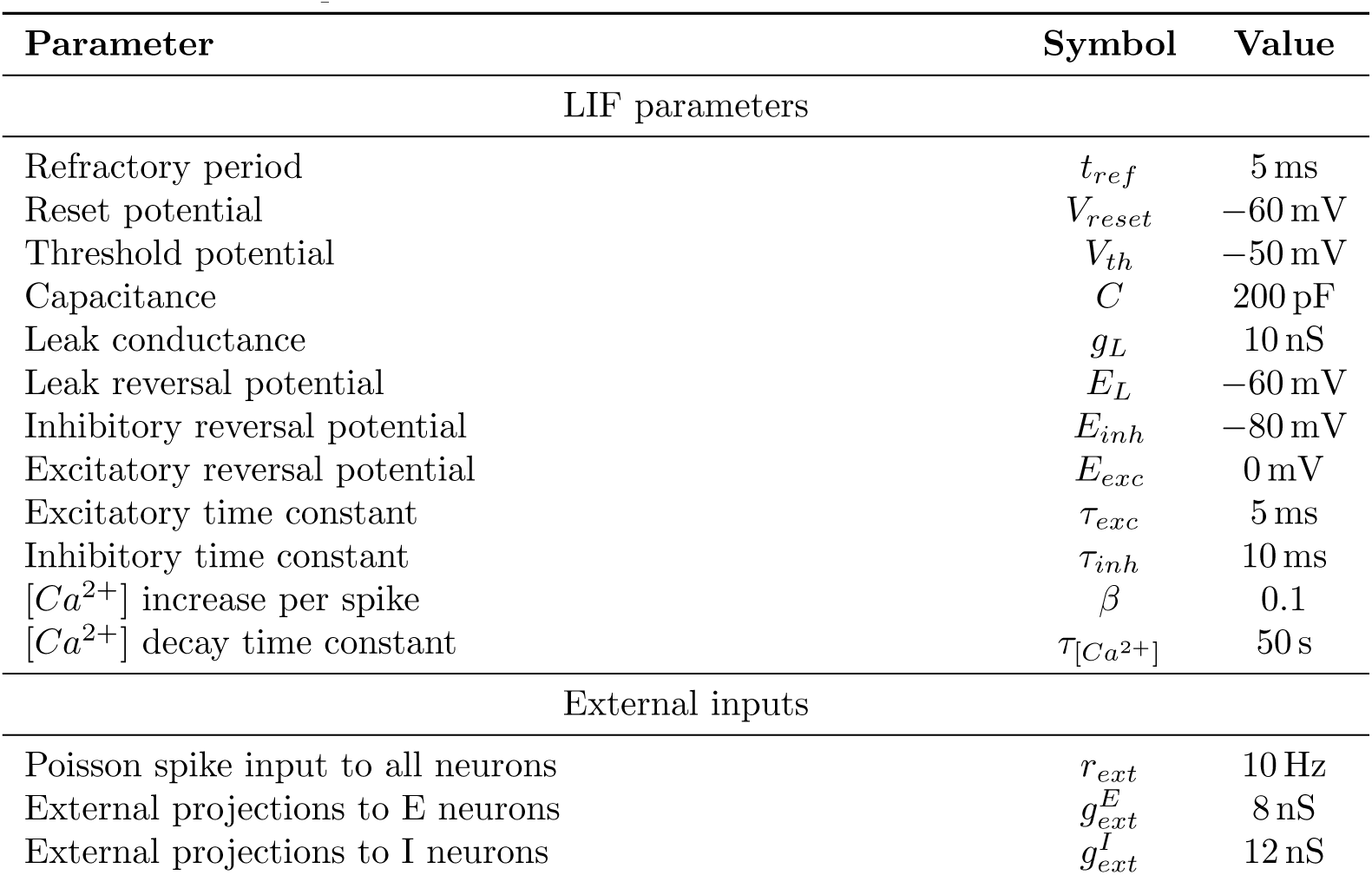
Neuronal parameters

Here, *τ*_[*Ca*^2+^]_ is the time constant with which the [*Ca*^2+^] decays in the absence of a spike, and *β* is the constant increase in [*Ca*^2+^] caused by each spike. Based on evidence that the outgrowth of synaptic structures depends on the concentration of intracellular calcium in neurons [69, 70], the rate of change of each type of synaptic element, (*dz/dt*) is given by:

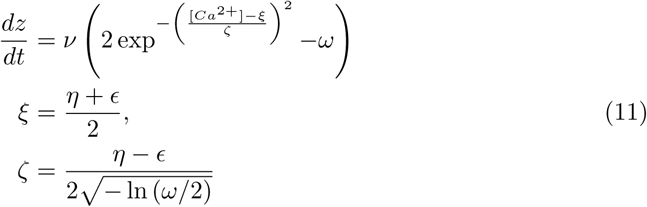

Here, *ν* is a scaling factor, *ξ* and *ζ* define the width and location of the Gaussian curve on the x-axis, while *ω* controls the location of the curve on the y-axis (0 < *ν*, 0 < *η* < *ϵ*, 0 < *ω* < 2). Given that ([*Ca*^2+^] *>* 0), (*dz/dt*) is bound as:

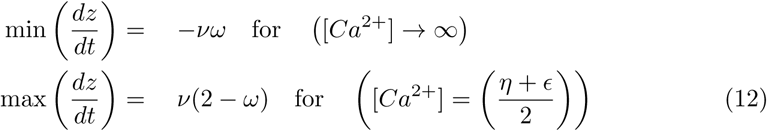

Within these bounds, as shown in Fig. 2, (*dz/dt*) is:

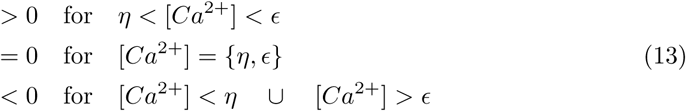

If, based on its activity, a neuron has more synaptic elements of a particular type (*z*) than are currently engaged in synapses (*z*_*connected*_), the free elements (*z*_*free*_) can participate in the formation of new synapses at the next connectivity update step:

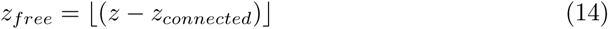

However, if they remain unconnected, they decay at each integration time step with a constant rate *τ*_*free*_:

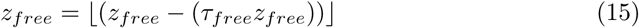

On the other hand, a neuron will lose *z*_*loss*_ synaptic connections if the number of a synaptic element type calculated by the growth rules (*z*) is less than the number of connected synaptic elements of the same type (*z*_*connected*_):

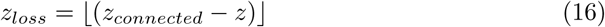

Table 5 lists the parameters governing the growth rules for all neurites.

**Table 5.** Growth rule parameters

| Parameter | Symbol | Value |
| --- | --- | --- |
| Optimal [$Ca^{2+}$] | $\psi$ | |
| Excitatory neurons |  |  |
| Scaling factor: pre-synaptic structures ($z_{pre}^E$) | $\nu_{pre}^E$ | $15 \times 10^{-4}$ |
| Vertical shift | $\omega_{pre}^E$ | $1 \times 10^{-2}$ |
| X-axis parameters | $(\eta_{pre}^E, \epsilon_{pre}^E)$ | $(\psi, 1.75 \times \psi)$ |
| Decay rate | $\tau_{pre, free}^E$ | 0.01 |
| Scaling factor: excitatory post-synaptic structures ($z_{post, E}^E$) | $\nu_{post, E}^E$ | $3 \times 10^{-5}$ |
| Vertical shift | $\omega_{post, E}^E$ | $4 \times 10^{-1}$ |
| X-axis parameters | $(\eta_{post, E}^E, \epsilon_{post, E}^E)$ | $(0.25 \times \psi, \psi)$ |
| Decay rate | $\tau_{post, E, free}^E$ | 0.01 |
| Scaling factor: inhibitory post-synaptic structures ($z_{post, I}^E$) | $\nu_{post, I}^E$ | $3 \times 10^{-4}$ |
| Vertical shift | $\omega_{post, I}^E$ | $4 \times 10^{-2}$ |
| X-axis parameters | $(\eta_{post, I}^E, \epsilon_{post, I}^E)$ | $(\psi, 3.5 \times \psi)$ |
| Decay rate | $\tau_{post, I, free}^E$ | 0.01 |
| Inhibitory neurons |  |  |
| Scaling factor: pre-synaptic structures ($z_{pre}^I$) | $\nu_{pre}^I$ | $3 \times 10^{-2}$ |
| Vertical shift | $\omega_{pre}^I$ | $4 \times 10^{-4}$ |
| X-axis parameters | $(\eta_{pre}^I, \epsilon_{pre}^I)$ | $(0.25 \times \psi, \psi)$ |
| Decay rate | $\tau_{pre, free}^I$ | 0.01 |
| Scaling factor: excitatory post-synaptic structures ($z_{post, E}^I$) | $\nu_{post, E}^I$ | $3 \times 10^{-5}$ |
| Vertical shift | $\omega_{post, E}^I$ | $4 \times 10^{-1}$ |
| X-axis parameters | $(\eta_{post, E}^I, \epsilon_{post, E}^I)$ | $(0.25 \times \psi, \psi)$ |
| Decay rate | $\tau_{post, E, free}^I$ | 0.01 |
| Scaling factor: inhibitory post-synaptic structures ($z_{post, I}^I$) | $\nu_{post, I}^I$ | $3 \times 10^{-5}$ |
| Vertical shift | $\omega_{post, I}^I$ | $4 \times 10^{-1}$ |
| X-axis parameters | $(\eta_{post, I}^I, \epsilon_{post, I}^I)$ | $(\psi, 3.5 \times \psi)$ |
| Decay rate | $\tau_{post, I, free}^I$ | 0.01 |

### Network simulations

Our network model is derived from the cortical network model proposed by Vogels et al. [41] that is balanced by inhibitory homeostatic STDP. Like the cortex, this network model is characterised by low frequency AI firing of neurons. Additionally, this network model has also been demonstrated to store attractorless associative memories for later recall. The simulation is divided into multiple phases, as shown in Fig. 14. These are documented in the following sections in detail.

**Fig 14.**
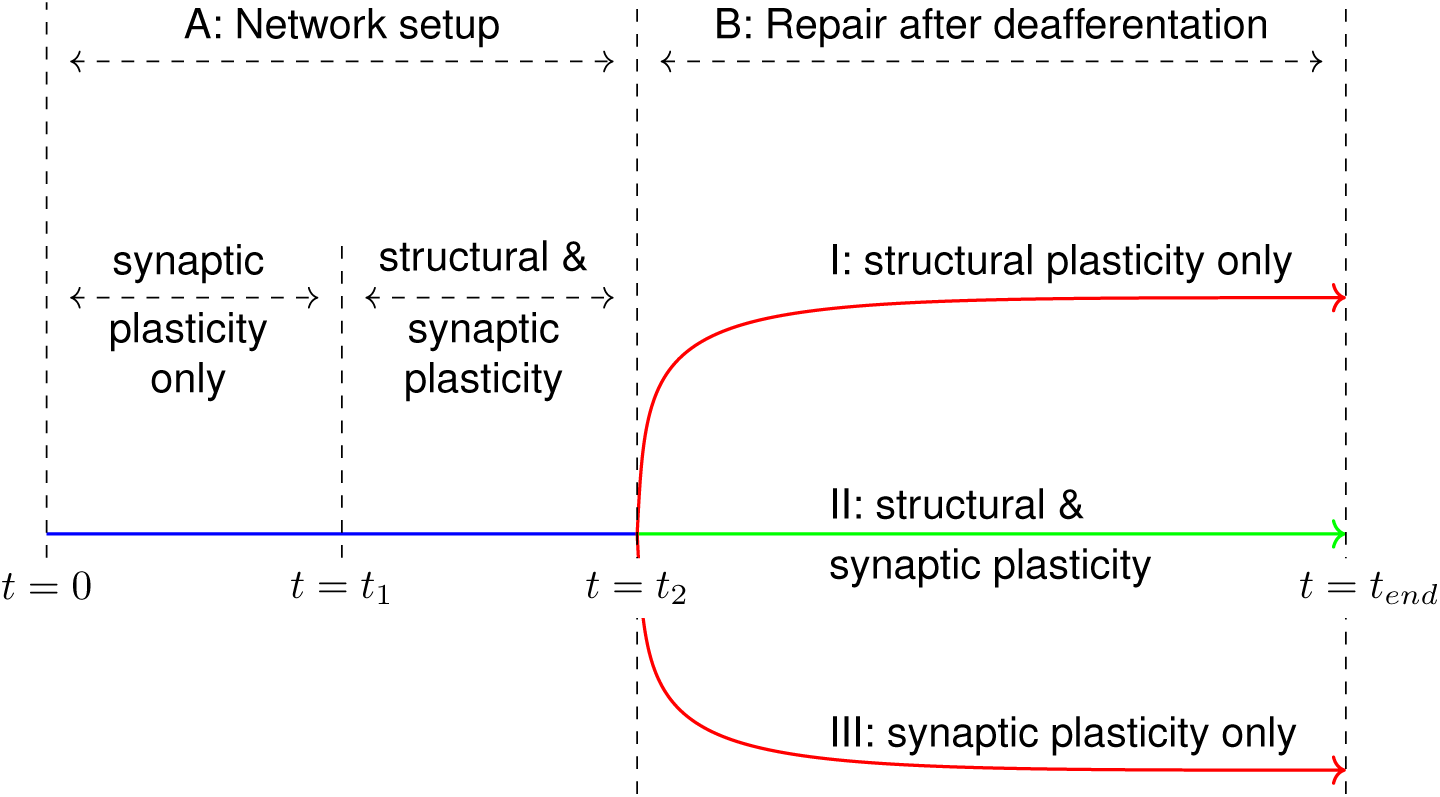
The simulation runs in 2 phases. Initially, the setup phase (0 s < *t* < *t*_2_) is run to set the network up to the balanced AI state. At (*t* = *t*_2_), a subset of the neuronal population is deafferented to simulate a peripheral lesion and the network is allowed to organise under the action of homeostatic mechanisms until the end of the simulation at (*t* = *t*_*end*_). Each homeostatic mechanism can be enabled in a subset of neurons to analyse its effects on the network after deafferentation.

#### Initial network structure

We simulate a network of *N*_*E*_ excitatory and *N*_*I*_ inhibitory neurons (*N*_*E*_/*N*_*I*_ = 4). Excitatory neurons are distributed in a two-dimensional rectangular plane such that the distance between two adjacent excitatory neurons is 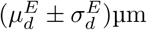. Inhibitory neurons are scattered such that they are evenly dispersed among the excitatory neurons such that the mean distance between adjacent inhibitory neurons is 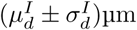. The rectangular plane is wrapped around as a toroid to prevent any edge effects from affecting the simulation. Table 6 summarises the parameters used to arrange the neurons.

**Table 6.** Network simulation parameters

| Parameter | Symbol | Value |
| --- | --- | --- |
| Simulation parameters |  |  |
| Integration time step | $dt$ | 0.1 s |
| Structural plasticity update interval |  | 1 s |
| Network parameters |  |  |
| Number of E neurons | $N_E$ | 8000 |
| Number of I neurons | $N_I$ | 2000 |
| Dimension of 2D E neuron lattice | | $100 \times 80$ |
| Dimension of 2D I neuron lattice | | $50 \times 40$ |
| Mean distance between E neurons | $\mu_d^E$ | 150 $\mu\text{m}$ |
| STD distance between E neurons | $\sigma_d^E$ | 15 $\mu\text{m}$ |
| Mean distance between I neurons | $\mu_d^I$ | 300 $\mu\text{m}$ |
| STD distance between I neurons | $\sigma_d^I$ | 15 $\mu\text{m}$ |
| Neurons in LPZ C |  | 2.5 % |
| Neurons in LPZ B |  | 2.5 % |
| Neurons in P LPZ |  | 5 % |
| Remaining neurons |  | 90 % |
| Initial network sparsity | $p$ | 0.02 |
| Initial out-degree | $n_{out}$ | $p \times \text{total possible targets}$ |
| Simulation stages |  |  |
| Synaptic plasticity only |  | 1500 s |
| Synaptic and structural plasticity |  | 500 s |
| Network deafferented at |  | 2000 s |

At (*t* = 0 s in Fig. 14), neurons in the network are connected such that the network has a sparsity of *p*. For each neuron, *n*_*out*_ targets are chosen from the complete set of possible post-synaptic neurons in a distance dependence manner as summarised in previous sections. Initially, static synapses in the network (II, IE, EI) are initialised to their mean conductances. The plastic (IE) synapses are subject to the homeostatic inhibitory synaptic plasticity mediated STDP rule proposed by Vogels, Sprekeler et al. [41] and are initialised to zero conductances.

External input to each neuron is modelled as an independent Poisson spike train with a mean firing rate *r*_*ext*_. These spike trains project on to excitatory and inhibitory neurons via static excitatory synapses with conductances 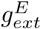 and 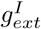 respectively. Figure 1A shows the various sets of synapses in the network.

#### Initial stabilisation to physiological state

The simulation is then started and the network permitted to stabilise to its balanced AI state until (*t* = *t*_2_ in Fig. 14). This phase consists of two simulation regimes. Initially, only inhibitory synaptic plasticity is activated to stabilise the network (*t* < *t*_1_ in Fig. 14).

As this state (*t* = *t*_2_ in Fig. 14) is considered the normal physiological state of our network model, the network parameters obtained at this point are set as the steady state parameters of neurons and synapses in the network. The optimal activity of each neuron, *ψ*, is set to the activity achieved by the neuron at this point, and its growth curves are initialised in relation to it. The mean conductance for new IE synapses is also set as the mean conductance of the IE synapses obtained at this stage.

Our implementation of homeostatic structural plasticity is then activated in the network at this point (*t* = *t*_1_ in Fig. 14) to verify that the network continues to remain in its balanced AI state in the presence of both homeostatic mechanisms.

#### Simulation of peripheral lesion

Next at (*t* = *t*_2_ in Fig. 14), the external Poisson spike train inputs are disconnected from excitatory and inhibitory neurons that fall in the LPZ to simulate a peripheral lesion in the network. For analysis, the neuronal plane is classified into four regions:

- LPZ C: the centre of the LPZ (Red in Fig. 1B).
- LPZ B: the inner border of the LPZ (Yellow in Fig. 1B).
- P LPZ: peri-LPZ, the outer border of the LPZ (Green in Fig. 1B).
- Other neurons: neurons further away from the LPZ (Grey in Fig. 1B).

#### Network reorganisation

The deafferented network is permitted to reorganise itself under the action of the active homeostatic mechanisms until the end of the simulation (*t* = *t*_*end*_ in Fig. 14). By selectively activating the two homeostatic mechanisms in different simulation runs, we were also able to investigate their effects on the network in isolation.

##### Structural plasticity mediated connectivity updates

All synapses in the network, except the connections that project the external stimulus on to the neuronal population, are subject to structural plasticity (Fig. 1A).

Free excitatory pre-synaptic and excitatory post-synaptic elements can combine to form excitatory synapses (EE, EI). Analogously, inhibitory pre-synaptic and inhibitory post-synaptic elements can plug together to form inhibitory synapses (II, IE). The set of possible partners for a neuron, therefore, comprises of all other neurons in the network that have free synaptic elements of the required type. From this set, *z*_*free*_ partners are chosen based on a probability of formation, *p*_*form*_, which is a Gaussian function of the distance between the pair, *d*:

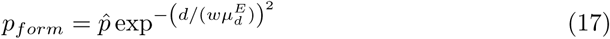

Here, 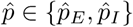 is the maximum probability, 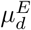 is the mean distance between two adjacent excitatory neurons, and *w* ∈ *w*_*E*_, *w*_*I*_ is a multiplier that controls the spatial extent of new synaptic connections.

Investigations indicate that lateral connections in the primary visual cortex are organised in a “Mexican hat” pattern. While experimental work does support the presence of the “Mexican hat” pattern [71, 72], anatomical research suggests that inhibitory connections are more localised than excitatory ones, contradicting the traditional use of shorter excitatory and longer inhibitory connections in computer models [73]. Analysis of the local cortical circuit of the primary visual cortex suggests that the “Mexican hat” pattern can either be generated by narrow but fast inhibition, or broad and slower inhibition that may be provided by longer axons of GABAergic basket cells [74, 75]. Investigations into the maintenance of the “Mexican hat” pattern are beyond the scope of this study. We therefore, limit ourselves to the traditional model of longer inhibitory connections and shorter local excitatory connections in this work by using a larger multiplier for inhibitory synapses, *w*_*I*_, than for excitatory synapses, *w*_*E*_, (*w*_*E*_ < *w*_*I*_).

New synapses that are added to the network are initialised with conductances similar to that of existing synapses in the balanced network. Their conductance values are taken from a Gaussian distribution centred at the mean conductance for that synapse type. Since new synapses can, therefore, be weaker or stronger than existing ones, this prevents the same set of synapses from being modified in each connectivity update.

In spite of them being plastic, the same method is also used for IE synapses. IE synapses are initialised with zero conductances at the start of the simulation and modify their strengths based on STDP [41]. When the network has achieved the balanced AI state, these conductances also settle at higher values. If new IE synapses formed after this point by structural plasticity were to be initialised to zero conductances, they would most likely be selected for deletion repeatedly as the weakest ones. STDP does not modulate inactive synapses either—synapses between pairs of neurons that have both been rendered inactive by deafferentation will not be weakened, and may not be lost. Therefore, to ensure the turnover of a diverse set of IE synapses also, new connections of this type are supplied with conductances similar to that of existing stable IE synapses in the balanced network.

Experimental evidence suggests that the stability of synapses is proportional to their efficacy [13, 76]. Taking this into account, we calculate the probability of deletion of a synapse, *p*_*del*_, as a function of its conductance *g*:

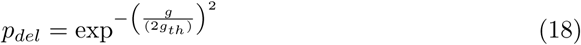

Here, *g*_*th*_ is a threshold conductance value calculated during the simulation, synapses stronger than which are considered immune to activity dependent changes in stability. They are removed from the list of options from which *z*_*loss*_ synapses are selected for deletion and are therefore, not considered for deletion at all.

For simplicity, for static excitatory synapses that all have similar conductances (EI, EE), we do not use this method of deletion. Instead, for these, *z*_*loss*_ connections are randomly selected for deletion from the set of available candidates. While II synapses are also static, the deletion of an inhibitory synapse by the loss of an inhibitory post-synaptic element can occur by the removal of either an IE or an II synapse. Therefore, to permit competition between II and IE synapses for removal, we apply weight based deletion to both these synapse sets.

The numbers of synaptic elements are updated at every simulator integration time step internally in NEST. Connectivity updates to the network, however, require updates to internal NEST data structures and can only be made when the simulation is paused. This increases the computational cost of the simulation, and we only make these updates at 1 s intervals. Gathering data on conductances, connectivity, and neuronal variables like [*Ca*^2+^] also require explicit NEST function calls while the simulation is paused. Therefore, we also limit dumping the required data to files to regular intervals. Table 7 summarises the various synaptic parameters used in the simulation.

**Table 7.**
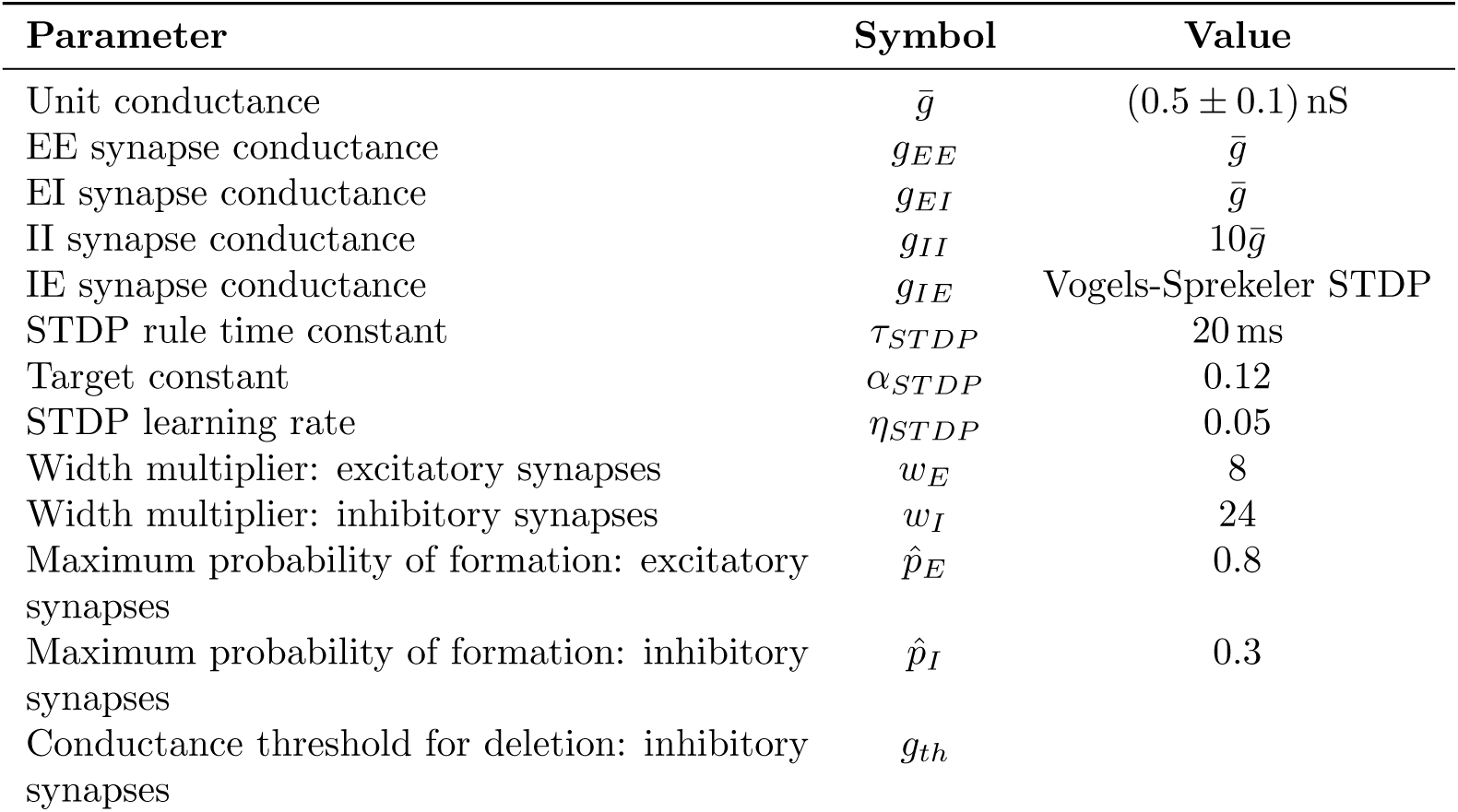
Synapse parameters

### Single cell simulations

We also studied the effects of our structural plasticity hypotheses in individual neurons using single neuron simulations. Figure 10A shows a schematic of our single neuron simulations.

The neuron is initialised to a steady state where it exhibits an indegree similar to neurons in the network simulations when in their AI state. To do so, a constant baseline input current *I*_*ext*_ is supplied to the neuron to provide it with activity. The [*Ca*^2+^] obtained by the neuron at this time is assumed as its optimal level, *ψ*. Using identical values of *η* and but different *ν* values for excitatory and inhibitory post-synaptic elements (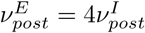 to mimic the initial indegree of neurons in our network simulations), and an input current that deviates the activity of the neuron off its optimal level (< *I*_*ext*_), the neuron is made to sprout 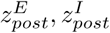 excitatory and inhibitory post-synaptic elements respectively 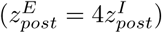. By assuming that each dendritic element receives inputs via conductances as observed in network simulations (*g*_*EE*_, *g*_*IE*_), the net input to the neuron that results in its activity can be approximated as:

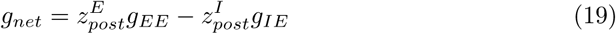

At this stage, the neuron resembles a one in network simulations in its balanced state before deafferentation. The current input is returned to its baseline value, thus returning the [*Ca*^2+^] to its optimal value, *ψ*. In addition, the growth curves for the neuron are restored as per our activity dependent structural plasticity hypotheses to verify that the neuron does not undergo any structural changes at its optimal activity level.

The external current input to the neuron is modulated sinusoidally to fluctuate the neurons [*Ca*^2+^] (Fig. 10B), and resultant changes in the numbers of its post-synaptic elements are recorded. As the neuron modifies its neurites, the change in excitatory and inhibitory input conductance received as a result is calculated (Fig. 10C).

## Acknowledgments

We are grateful to Benjamin-Torben Nielsen for fruitful discussions and feedback on the work.

This work has made use of the University of Hertfordshire’s high-performance computing facility.

We are also most grateful to the NEST development team, in particular to Sandra Diaz-Pier, for discussions and assistance with the modelling of structural plasticity in the NEST simulator.

## Notes

### Competing Interest Statement

The authors have declared no competing interest.

### Summary of Updates

Figure 11 was added to improve the section on the "activity dependent dynamics of pre-synaptic structures".

